# High-fructose feeding suppresses cold-stimulated brown adipose tissue glucose uptake in young men independently of changes in thermogenesis and the gut microbiome

**DOI:** 10.1101/2022.01.11.475847

**Authors:** Gabriel Richard, Denis P. Blondin, Saad A. Syed, Laura Rossi, Michelle E. Fontes, Mélanie Fortin, Serge Phoenix, Frédérique Frisch, Stéphanie Dubreuil, Brigitte Guérin, Éric E. Turcotte, Martin Lepage, Michael G. Surette, Jonathan D. Schertzer, Gregory R. Steinberg, Katherine M. Morrison, André C. Carpentier

## Abstract

Diets rich in added sugars, especially high in fructose, are associated with metabolic diseases such as insulin resistance, and non-alcoholic fatty liver disease. Studies have shown a link between these pathologies and changes in the microbiome and its metabolites. Given the reported associations in animal models between the microbiome and brown or beige adipose tissue (BAT) function, and the alterations in the microbiome induced by high glucose or high fructose diets, we investigated the potential causal link between high glucose or fructose diets and BAT dysfunction in humans. We show that BAT glucose uptake, but not thermogenesis, is impaired by a high fructose but not high glucose diet, in the absence of changes in body mass, the gastrointestinal microbiome, and faecal short-chain fatty acids. We conclude that BAT metabolic dysfunction occurs independently from changes in gut microbiome composition, and earlier than other pathophysiological abnormalities associated with insulin resistance and dyslipidemia during fructose overconsumption in humans.

## Introduction

The consumption of fructose, a food additive highly prevalent in ultra-processed foods in the form of sucrose or high-fructose corn syrup, has risen precipitously over the past few decades. Its overconsumption has paralleled the rapid rise in obesity and the associated metabolic complications including insulin resistance, non-alcoholic fatty liver disease (NAFLD) and cardiovascular disease (CVD). Animal studies, primarily performed in rodents, have consistently demonstrated that diets high in sucrose or fructose induce hypertriglyceridemia and hyperinsulinemia, providing a causal link between fructose consumption and dysmetabolism (Hwang et al., 1987; Martinez et al., 1994; Nikkilä and Ojala, 1965; Zavaroni et al., 1982). Such direct evidence in humans, however, has remained more elusive. Several pivotal short-term intervention studies in humans have shown that consuming fructose-sweetened beverages contributing 10-25% of energy intake increases circulating triglyceride levels, visceral adiposity, hepatic lipid accretion, and insulin resistance compared to isocaloric glucose or sweeteners (Hieronimus et al., 2020; Schwarz et al., 2015; Stanhope et al., 2009, 2011, 2015). Conversely, restricting sugar-sweetened beverages in children with obesity (Lustig et al., 2016; Schwarz et al., 2017) lowered fasting triglyceride levels and hepatic fat, while also improving postprandial insulin kinetics. While these studies have been critical to the characterization of the impact of consuming modest levels of fructose on metabolic risk factors, the mechanisms by which isocaloric fructose may promote metabolic disease and its association with insulin resistance remain unknown. This has been particularly difficult to ascertain given the limited number of studies using appropriate controls and directly comparing the effects of fructose-sweetened beverages to isocaloric glucose.

One emerging mechanism that has been proposed as a driving factor in fructose-induced dysmetabolism is the progressive impairment in brown adipose tissue (BAT) metabolic activity. Retrospective studies commonly report that high spontaneous BAT activity is associated with a favorable metabolic profile including lower circulating triglycerides and higher insulin sensitivity (Becher et al., 2021; Cypess et al., 2009; Ouellet et al., 2011; Wibmer et al., 2021), whereas impairments in BAT glucose uptake, NEFA uptake, blood flow and insulin sensitivity have been noted in aging, obesity and type 2 diabetes (T2D) (Blondin et al., 2015a; Orava et al., 2013; Ouellet et al., 2011; Saari et al., 2020). Although several recent studies have demonstrated the ability to activate human BAT through acute and intermittent cold exposure and acute and chronic beta-adrenergic stimulation (Baskin et al., 2018; Blondin et al., 2014, 2020; Cypess et al., 2015; Hanssen et al., 2015, 2016; O’Mara et al., 2020), our understanding of the factors that suppress its activity with aging, obesity or T2D remains incomplete. Recent evidence in rodents suggests that fructose ingestion increases intestinal microbiota-derived acetate production, which may not only provide a precursor for hepatic lipogenesis (Zhao et al., 2020) but could incidentally suppress BAT function (Sun et al., 2021), providing a prospective causal role for BAT in fructose-associated dysmetabolism.

We hypothesized that fructose can impair BAT metabolic activity and thermogenesis and may be a causal factor to the fructose-induced deterioration in metabolic profile in humans. To test this hypothesis, we conducted a randomized crossover 2-week fructose *vs.* glucose overfeeding (+25% of energy intake) study in 10 young healthy men interspersed by a 4-week washout period and preceded by a metabolic assessment performed under an isocaloric condition. The primary outcomes compared microbiome composition, faecal short-chain fatty acids (SCFA), BAT oxidative metabolism and BAT intracellular triglyceride content between each 2-week diet. Whole-body energy metabolism and BAT glucose metabolism were also investigated as secondary outcomes. A complete list of the primary and secondary outcomes for this study is available on clinicaltrials.gov (NCT03188835).

## Results

### Short-term fructose overfeeding does not alter adiposity, the gut microbiome or faecal short-chain fatty acids

The high fructose and high glucose diets were designed to be hypercaloric with the carbohydrate-enriched drink representing an additional 25 % of the total caloric intake. Participants confirmed drinking the entire diet supplement and were instructed to keep their other dietary intake the same as in the baseline isocaloric period. Changes in total body mass, fat mass and lean mass (after the diet versus before the diet) were highly variable between individuals for all diets (Figure 1 A). On average, there was no difference in body composition between diets. The high fructose diet elicited a more reproducible increase in total body weight, although this seemed to be related to a gain in lean body mass, rather than in fat mass. Physical activity levels and resting energy expenditure did not change between diets (Figure 1 B–C). Similarly, there were no differences in tissue-specific fat content assessed by MRI for BAT, subcutaneous or visceral adipose tissue, or the liver (Figure 1 D).

**Figure 1.**
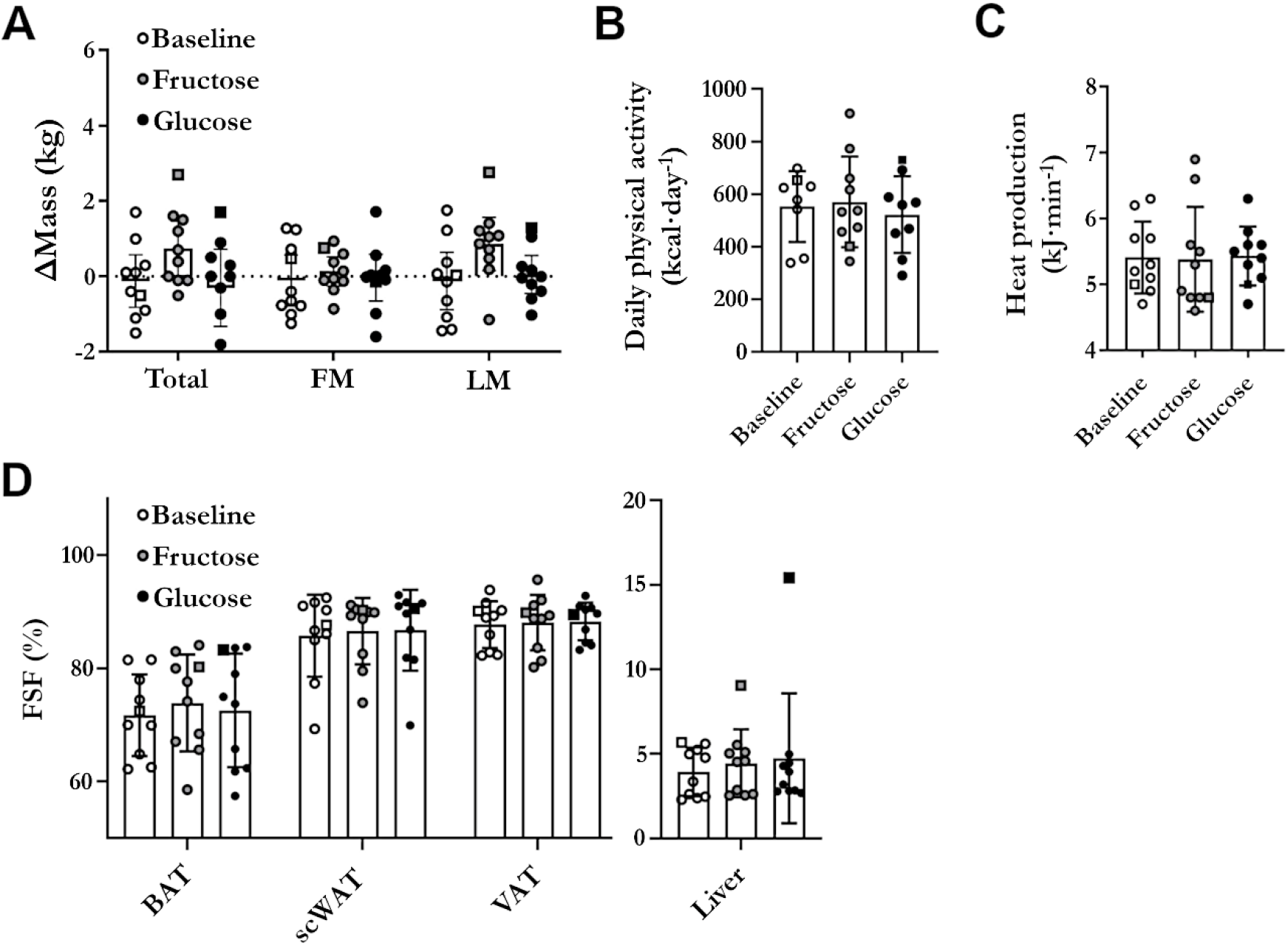
Body composition at baseline and following the high-carbohydrate diets. No significant change in whole-body or tissue specific composition was noted with the different diets. (A) Changes in body mass and composition assessed (after the diet - before the diet) by dual-energy X-ray absorptiometry (n = 10). (B) Energy expenditure from exercise assessed by uniaxial accelerometer at baseline (n = 8), high fructose (n = 10), and high glucose (n = 9). (C) Whole-body energy expenditure at rest assessed by indirect calorimetry (n = 10). (D) Fat content of tissues assessed by Dixon MRI (n = 10). BAT: brown adipose tissue; FM: fat mass; FSF: fat signal fraction; LM: lean mass; scWAT: subcutaneous white adipose tissue; VAT: visceral adipose tissue. Participant 13 was excluded from the analyses due to antibiotic use (potential effects on the microbiome) before the fructose and glucose diets and is shown with a square symbol shape.

We investigated the effects of the high-carbohydrate diets on the faecal microbiome composition through 16S ribosomal RNA gene variable region 3-4 sequencing. Participants provided stool samples at the end of every 2-week diet period (at baseline, high glucose, and high fructose). No significant differences in faecal microbiome composition were observed in terms of taxonomic composition, α-, and β-diversity (Figure 2 A–C). We also measured the faecal SCFA content (acetate, butyrate, and propionate), important metabolites produced by microbiota fermentation of indigestible carbohydrates, by gas chromatography–tandem mass spectrometry. There was no effect of the diets on SCFA content or content ratios (e.g., acetate/propionate ratio), although there was a high variability between subjects (Figure 2 D–E).

**Figure 2.**
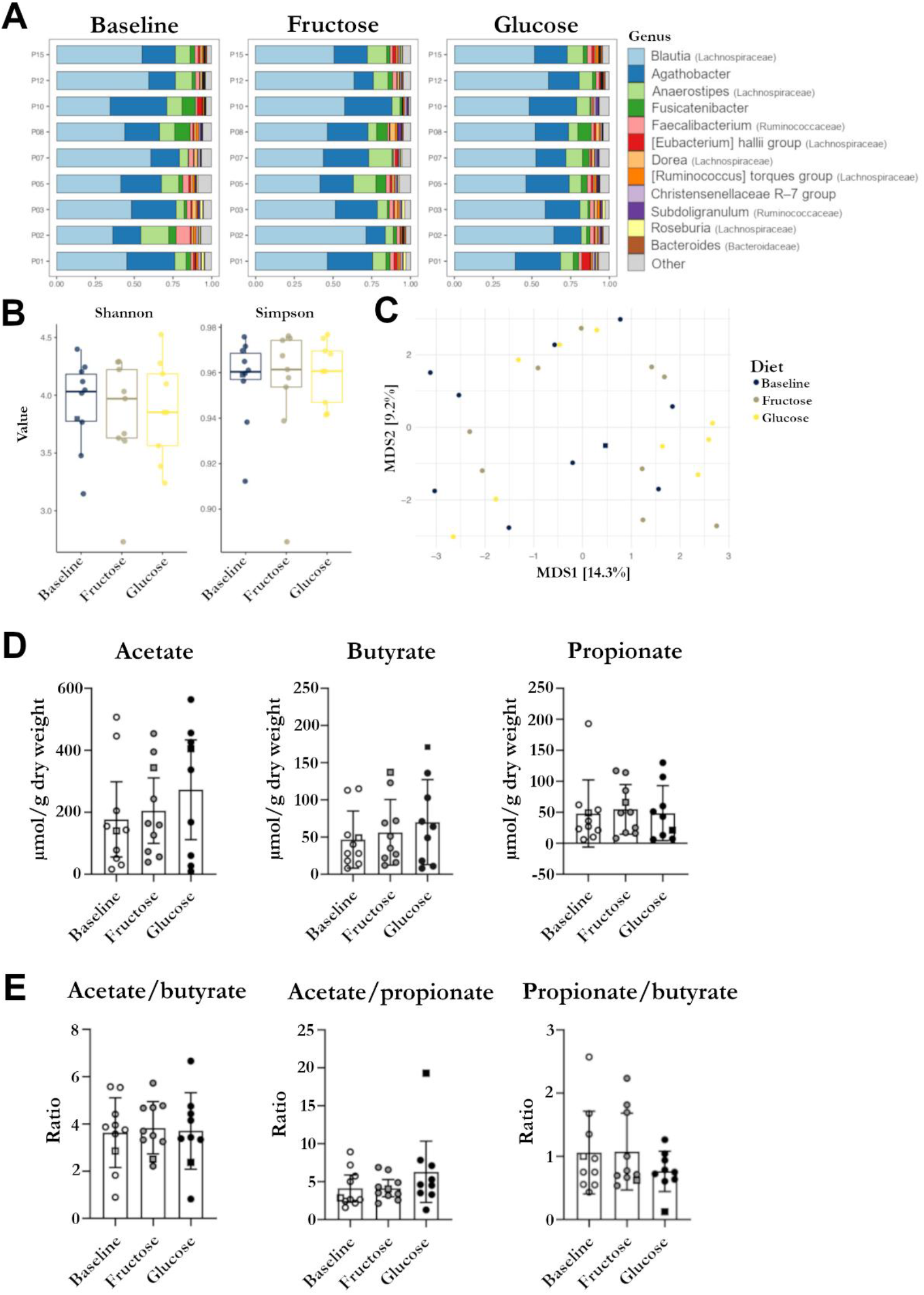
Fecal microbiome analyses and faecal short-chain fatty acids for the different diets. (A) Relative abundance taxonomic summaries divided by diet and ordered by participant for all participants included in statistical analyses (n = 9). Top 12 genera in the data were coloured and are identified in the legend. All other genera are in ‘other’. (B) Participants’ fecal microbial α-diversity assessed by Shannon and Simpson indices at baseline (n = 10), high fructose (n = 9) and high glucose (n = 9). Each dot represents the value of the diversity metric from each sample. A generalized mixed-effect model found no significant differences by α-diversity. (C) Participants’ fecal microbial β-diversity assessed by Aitchison distance and visualized using a Principal Coordinate Analysis (PCoA) plot. Each dot represents a sample and samples are colored by diet (baseline [n = 10], high fructose [n = 9] and high glucose [n = 9]). A PERMANOVA found no significant differences by β-diversity. Percent variation explained by each axis is noted in square brackets. (D) Fecal short-chain fatty acid composition for baseline (n = 10), high fructose (n = 10) and high glucose (n = 9). No significant difference was observed between diets. (E) Fecal short-chain fatty acid ratios. No significant difference was observed between diets. Participant 13 was excluded from the analyses due to antibiotic use (potential effects on the microbiome) before the fructose and glucose diets and is shown with a square symbol shape.

### Cold-induced BAT oxidative metabolism and intracellular triglycerides depletion remain normal after the carbohydrate-enriched diets

Participants underwent mild cold exposure (18 °C) to activate BAT metabolism (Ouellet et al., 2012). Primary outcomes were BAT oxidative metabolism assessed with ^11^C-acetate positron emission tomography (PET), as well as intracellular fatty acid content and composition assessed by magnetic resonance imaging and spectroscopy (MRI and MRS).

During the intervention, skin temperature, shivering activity, heat production, circulating energy substrates, whole-body substrate oxidation, and hormones were also monitored (Table 1). As expected, cold exposure elicited a decrease in mean skin temperature and increase in heat production, and this was fueled mainly by an increase in fatty acid oxidation. Both blood glucose and insulin levels decreased during cold exposure while non-esterified fatty acids increased. This is consistent with an increase in glucose uptake by tissues and an increase in whole-body adipose tissue intracellular lipolysis.

**Table 1.**
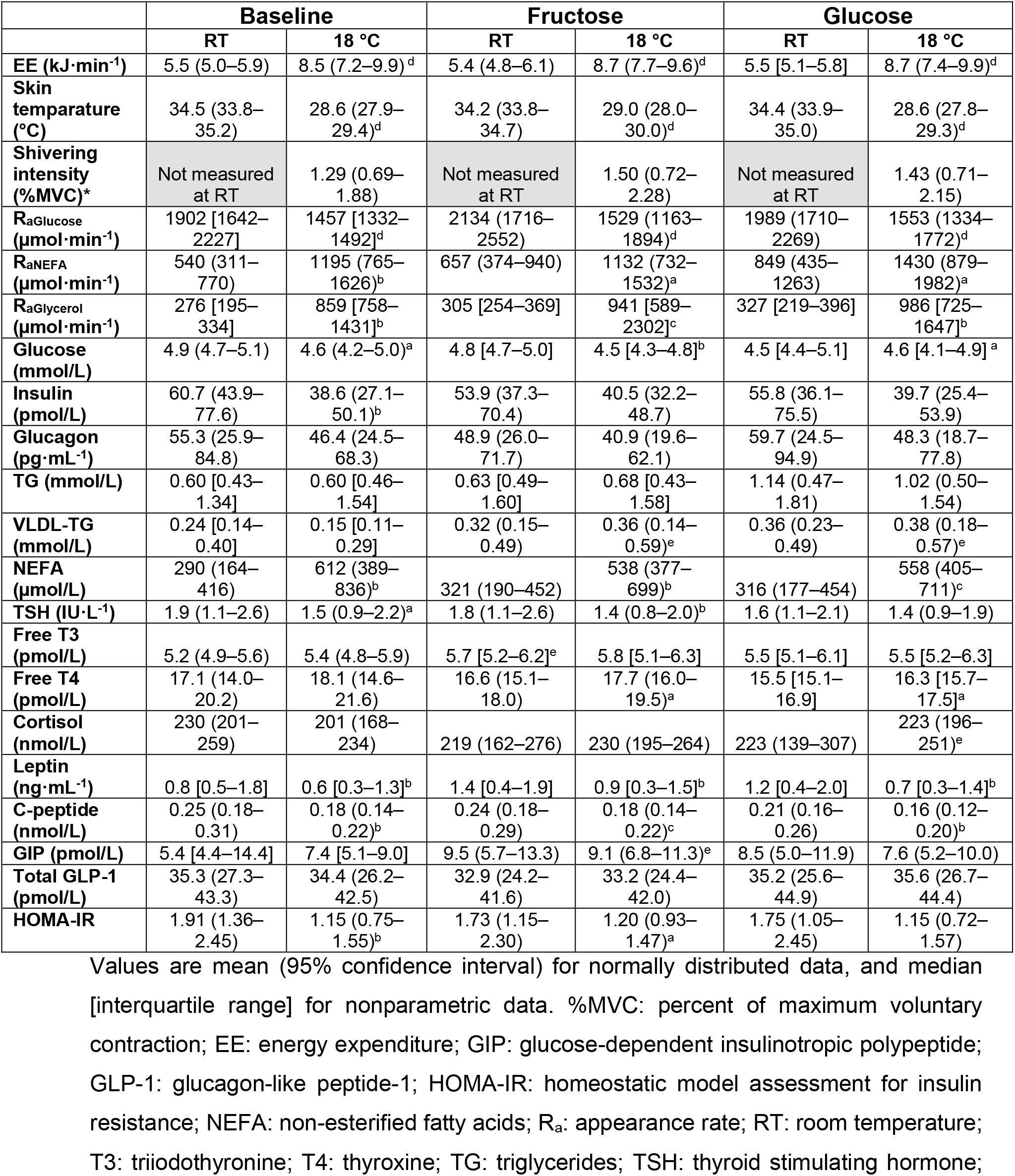
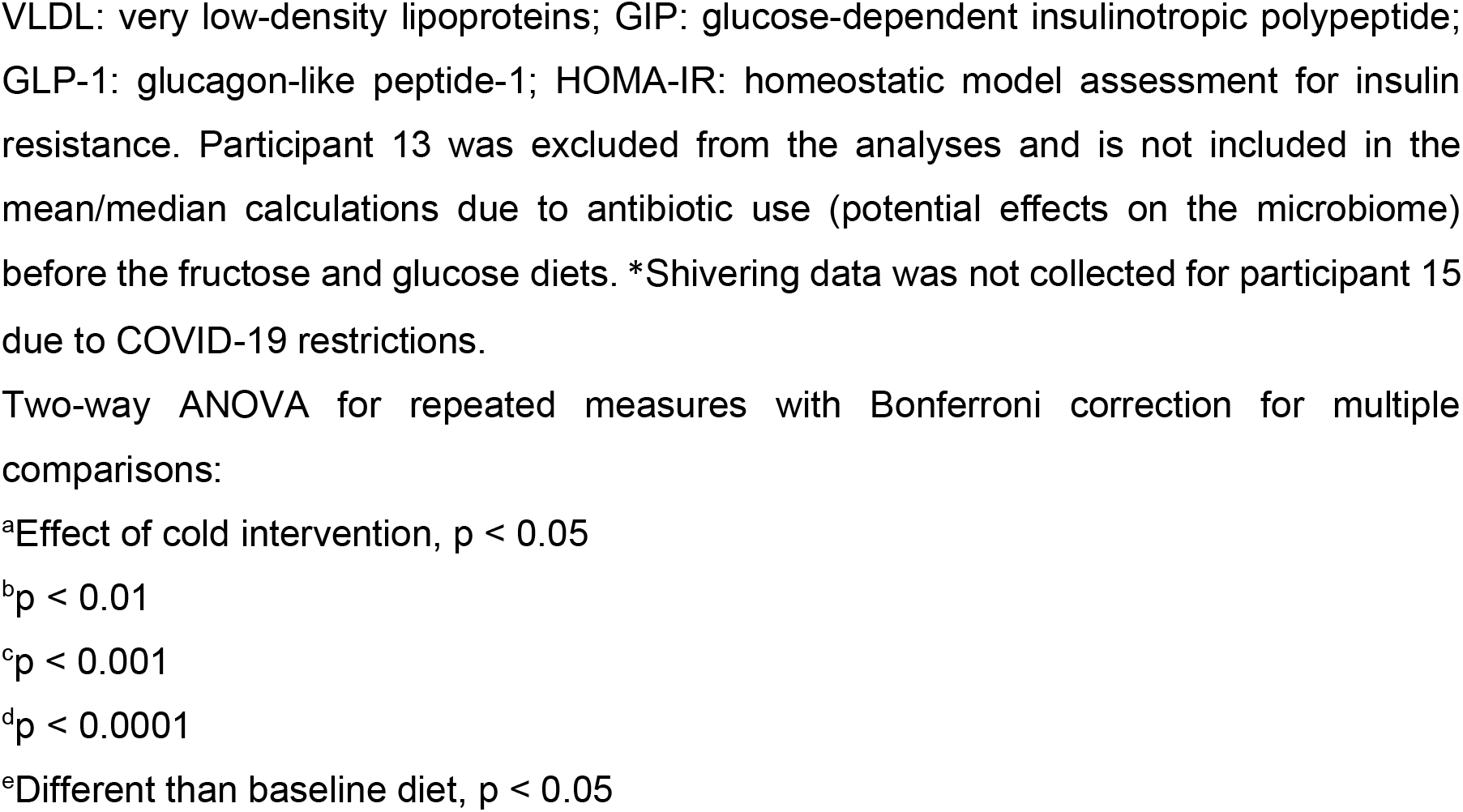
Whole-body response to diets and cold.

The time-activity curves of ^11^C-acetate in BAT showed a higher uptake and washout during cold exposure than at room temperature (Figure 3 A). These changes were quantified with a four-compartment, two-tissue model developed by our group (Richard et al., 2021, 2019). With this model, the uptake parameter (*K*1) can be used to estimate blood flow changes because first-pass extraction of the tracer is very high in most tissues (Labbé et al., 2011). The washout parameter (*k*2) represents oxidation (*i.e*., BAT thermogenesis). There was a significant increase in blood flow (Figure 3 B; p = 0.007) and oxidation (Figure 3 C; p = 0.003) during cold exposure consistent with increased mitochondrial respiration. However, there was no significant difference between diets in blood flow (fructose: p = 0.3; glucose p = 0.5) or oxidation (fructose and glucose: p = 0.6) during cold exposure, suggesting that mitochondrial respiration is not affected by 2 weeks of high carbohydrate diet. Measures at room temperature were only acquired during the baseline protocol to limit radiation exposure to participants. Therefore, the magnitude of change between room temperature and cold exposure is reported only at baseline.

**Figure 3.**
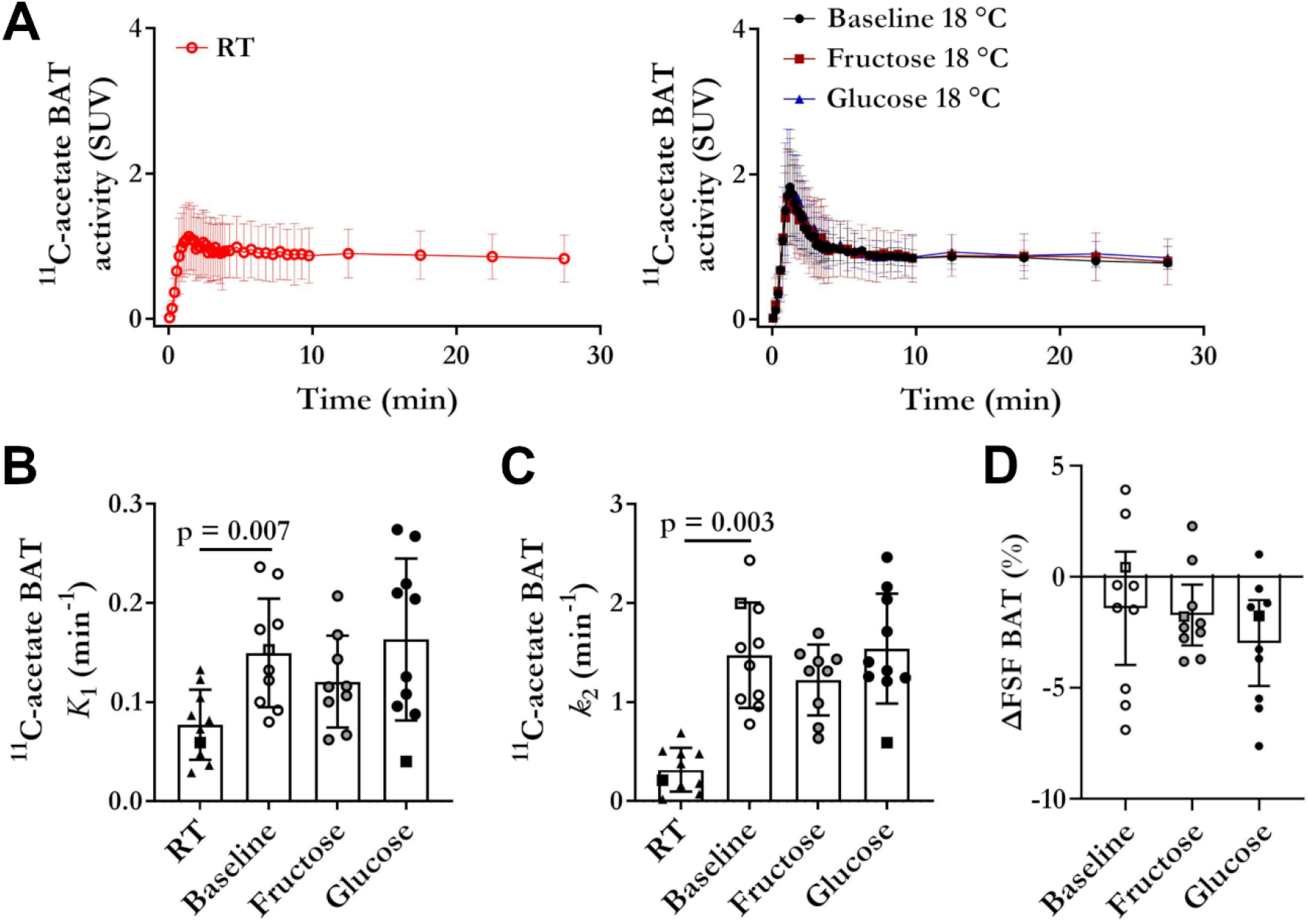
Brown adipose tissue oxidative metabolism and intracellular triglyceride depletion. (A) Mean time-activity curves showing the uptake and elimination of ^11^C-acetate or its metabolites at room temperature and during cold exposure (18 °C) for baseline, glucose, and fructose diets. (B) Blood flow index at baseline (n = 10; room temperature and cold), high fructose (cold: n = 9), and high glucose (cold: n = 10). A paired one-way ANOVA found a significant difference between room temperature and baseline. (C) Oxidative metabolism at baseline (n = 10; room temperature and cold), high fructose (cold: n = 9), and high glucose (cold: n = 10). A paired one-way ANOVA found a significant difference between room temperature and baseline cold exposure. (D) Fat signal fraction shift in brown adipose tissue with cold exposure (18 °C - room temperature) for the different diets (n = 10). There is no difference in shifts between diets. BAT: brown adipose tissue; FSF: fat signal fraction; *K*1: ^11^C-acetate uptake rate; *k*2: ^11^C-acetate oxidation rate; RT: room temperature; SUV: standardized uptake value. Participant 13 was excluded from the analyses due to antibiotic use (potential effects on the microbiome) before the fructose and glucose diets and is shown with a square symbol shape. ^11^C-acetate was not performed for participant 13 fructose due to problems with tracer synthesis.

Intracellular triglyceride content, as assessed by Dixon MRI fat signal fraction, decreased in BAT during cold exposure (Figure 3 D). The reduction in BAT fat signal fraction following cold exposure was consistent with previous studies (Abreu-Vieira et al., 2020; Ahmed et al., 2021; Coolbaugh et al., 2019; Oreskovich et al., 2019) and with the previously reported increase in BAT radiodensity (Baba et al., 2010; Blondin et al., 2017; Gifford et al., 2016; Lundström et al., 2015), which both reflect a progressive decrease in intracellular triglyceride content. This is also consistent with previous findings by our group indicating that intracellular triglycerides are the preferred energy source for BAT thermogenesis (Blondin et al., 2017). Diets did not modify the BAT fat signal fraction at room temperature (Figure 1 D) nor the shift with cold exposure (Figure 3 D). These findings are in line with the previous observations that diets did not alter body composition, and that BAT oxidative metabolism was preserved.

Furthermore, the lipid composition of BAT and subcutaneous white adipose tissue investigated by MRS (Table 2) were not affected by diets or cold exposure. However, as expected (Hamilton et al., 2011), BAT contained more saturated fatty acids than scWAT, demonstrated by a lower number of double bonds (median for all diets and temperatures: 3.5 [IQR: 3.0–3.8] *vs*. 4.1 [IQR: 3.4–4.5]; p = 0.002) and methylene-interrupted single bonds (median for all diets and temperatures: 1.4 [IQR: 1.0–2.0] *vs*. 3.3 [IQR: 2.6–3.9]; p < 0.0001).

**Table 2.** Fat composition of different tissues assessed by MRI and MRS.

|  | Baseline |  | Fructose |  | Glucose |  |
| --- | --- | --- | --- | --- | --- | --- |
|  | RT | 18 °C | RT | 18 °C | RT | 18 °C |
| **BAT FSF (%)** | 71.8 (66.6–76.9) | 70.3 (64.3–76.4) | 73.2 (66.5–79.9) | 71.5 (63.7–79.3)<sup>a</sup> | 71.4 (63.8–79.0) | 68.3 (60.0–76.6)<sup>a</sup> |
| **scWAT FSF (%)** | 87.2 [83.1–91.2] | 88.3 [81.4–91.2] | 89.4 [81.1–90.2] | 89.6 [80.9–90.9] | 89.7 [81.7–91.6] | 89.9 [82.9–90.5] |
| **VAT FSF (%)** | 87.8 (84.8–90.7) | 87.9 (85.5–90.3) | 87.9 (83.9–91.9) | 88.2 (85.2–91.1) | 88.2 (85.5–90.9) | 87.0 (83.2–90.8) |
| **Liver FSF (%)** | 4.1 [2.5–5.3] | 4.5 [2.4–5.3] | 4.5 [2.6–5.1] | 5.1 [2.8–6.1]<sup>a</sup> | 3.6 (2.9–4.2) | 3.5 (2.6–4.4) |
| **BAT ndb** | 3.6 (2.5–4.7) | 3.9 (3.1–4.6) | 3.7 (2.7–4.7) | 3.1 (2.6–3.6) | 3.6 (2.7–4.5) | 3.2 (2.8–3.7) |
| **scWAT ndb** | 4.1 (3.0–5.2) | 3.5 (2.7–4.2) | 3.9 (3.2–4.5) | 4.1 (3.4–4.8) | 4.2 (3.4–5.1) | 3.7 (3.0–4.5) |
| **BAT nmdb** | 2.5 [1.7–3.4] | 2.5 [2.2–3.2] | 2.2 [1.8–3.3] | 2.0 [0.4–2.8] | 3.2 (1.7–4.7) | 2.0 (1.4–2.6) |
| **scWAT nmdb** | 3.5 (2.3–4.6) | 3.2 (2.6–3.8) | 3.2 (2.0–4.3) | 2.9 (2.1–3.7) | 3.9 (2.9–5.0) | 2.4 (1.4–3.5) |

Values are mean (95% confidence interval) for normally distributed data, and median [interquartile range] for nonparametric data. BAT: brown adipose tissue; FSF: fat signal fraction; ndb: number of double bonds; nmdb: number of methylene-interrupted double bonds; RT: room temperature; scWAT: lower back subcutaneous white adipose tissue; VAT: visceral adipose tissue. Participant 13 was excluded from the analyses and is not included in the mean/median calculations due to antibiotic use (potential effects on the microbiome) before the fructose and glucose diets.
Two-way ANOVA for repeated measures with Bonferroni correction for multiple comparisons:
<sup>a</sup>Effect of cold intervention, $p < 0.05$

### Short term fructose overfeeding impairs BAT glucose metabolism

Tissue-specific glucose metabolism was assessed by ^18^F-FDG PET during cold exposure. The time-activity curves of ^18^F-FDG in BAT (Figure 4 A) show a quick initial increase in signal, corresponding to the first-pass extraction of ^18^F-FDG by the tissue. Afterwards, the signal continues to increase at a slower pace as ^18^F-FDG in the blood is taken up and trapped into BAT. The initial uptake phase appears similar for all diets. However, the subsequent trapping rate of ^18^F-FDG is slower after the high fructose diet. To quantify this phenomenon, BAT fractional glucose uptake was assessed with a Patlak graphical model (Patlak and Blasberg, 1983). The tissue metabolic rate of glucose was then calculated using fractional glucose extraction, glucose concentration, and a lumped constant of 1.14 for fat and 1.16 for muscle (Peltoniemi et al., 2000; Virtanen et al., 2001). Both the fractional extraction (Figure 4 B; p = 0.044) and metabolic rate of glucose (Figure 4 C; p = 0.036) decreased in BAT following the high fructose diet, but not following the high glucose diet. We did similar measures and modeling in different skeletal muscles as well as subcutaneous adipose tissue and conclude that this decrease in glucose uptake was specific to BAT (Figure 4 D). The reduction in cold-stimulated BAT glucose uptake following the high fructose diet was not associated with differences in blood glucose, insulin, HOMA-IR, glucagon, cortisol, c-peptide, GIP or GLP1. Furthermore, no difference in whole-body carbohydrate oxidation was observed with diets (Table 1).

**Figure 4.**
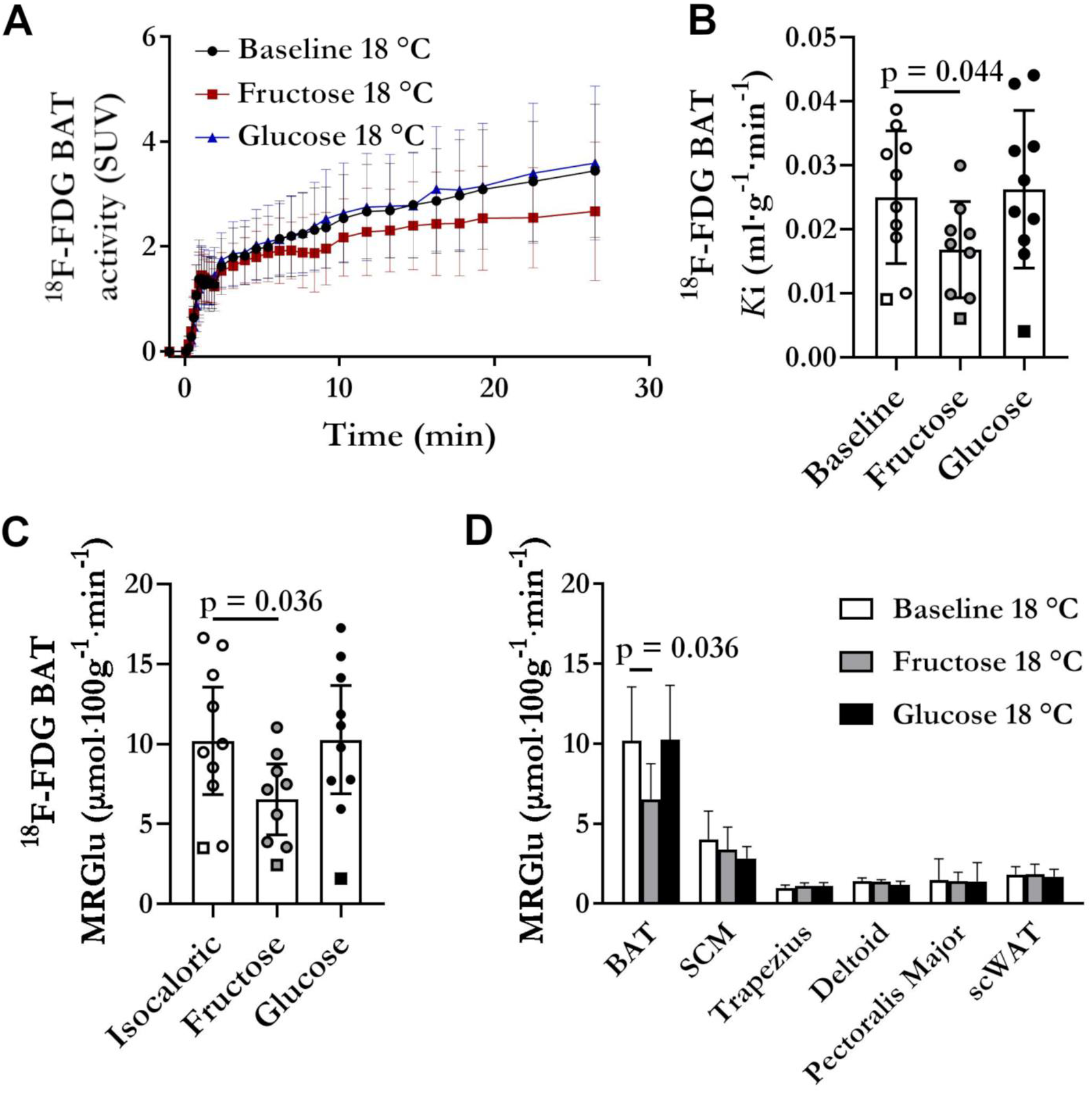
Glucose metabolism of tissues with cold exposure. (A) Mean brown adipose tissue time-activity curves showing ^18^F-fluorodeoxyglucose uptake during cold exposure (18 °C) for baseline, glucose, and fructose diets. (B) Brown adipose tissue fractional glucose uptake at baseline (cold: n = 10), high fructose (cold: n = 9), and high glucose (cold: n = 10). A paired one-way ANOVA found a significant difference between baseline and fructose. (C) Brown adipose tissue metabolic rate of glucose at baseline (cold: n = 10), high fructose (cold: n = 9), and high glucose (cold: n = 10). A paired one-way ANOVA found a significant difference between baseline and fructose. (D) Metabolic rate of glucose for brown adipose tissue compared to skeletal muscles and subcutaneous fat at baseline (cold: n = 10), high fructose (cold: n = 9), and high glucose (cold: n = 10). There was no significant difference for tissues other than BAT. ^18^F-FDG: ^18^F-fluorodeoxyglycose; BAT: brown adipose tissue; *K*i: fractional glucose uptake; MRGlu: metabolic rate of glucose; SCM: sternocleidomastoid muscle; scWAT: lower-back subcutaneous white adipose tissue; SUV: standardized uptake value. Participant 13 was excluded from the analyses due to antibiotic use (potential effects on the microbiome) before the fructose and glucose diets and is shown with a square symbol shape. Participant 2 fructose ^18^F-fluorodeoxyglucose was excluded due to excessive participant motion.

These observations point to a very localized impairment in glucose metabolism specific to BAT and independent of changes in oxidative metabolism or intracellular triglyceride depletion. Because ^18^F-FDG PET provides macroscopic data (with 4 mm^3^ image voxels) and does not discriminate between metabolic pools (i.e., ^18^F-FDG and ^18^F-FDG-6-phosphate produce the same signal), we cannot determine the exact cause of this impairment. Based on the ^11^C-acetate data, blood flow is not affected by the fructose diet.

Therefore, the reduction in glucose uptake is likely due to a deficit in transport into the brown adipocytes (e.g., insulin resistance), reduced phosphorylation rates or a combination of both. Whether this is related to a local effect of circulating acetate as suggested by Sun et al. (Sun et al., 2021) cannot be determined with our experimental design. There was no change in the acetate content of the stool samples, and plasma acetate could not be measured accurately by our gas chromatography-tandem mass spectrometry method.

### Fructose overfeeding increases liver triglyceride content as well as serum VLDL-TG in the cold

The liver triglyceride content assessed by MRI was significantly increased after cold exposure compared to room temperature only after the high fructose diet (Figure 5 A and Table 2; p = 0.037). This increase was not observed in the neighboring visceral or subcutaneous adipose tissues (Figure 5 A), and as noted previously, BAT triglyceride content decreased in the cold exposure condition (Figure 3 D). This suggests that fructose consumption promotes hepatic lipid storage under cold-induced increase in adipose tissue intracellular lipolysis, even though the magnitude of this increased lipolysis is not affected by diets (Figure 5 B–C).

**Figure 5.**
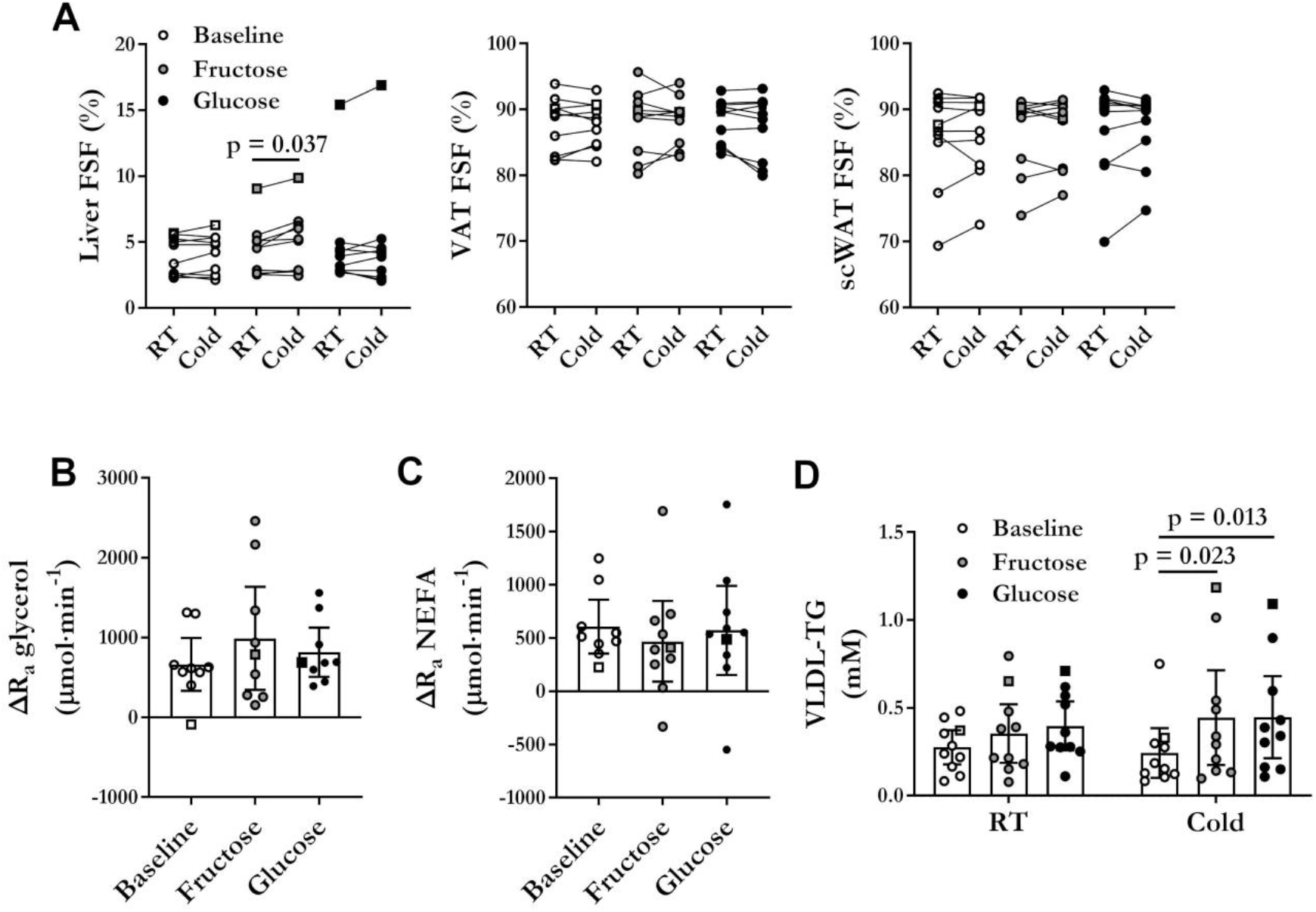
Effect of cold exposure on the liver and whole-body lipid metabolism. (A) Fat signal fraction of the liver, visceral adipose tissue, and white adipose tissue at room temperature and in the cold (18 °C) for each diet (n = 10). A two-way ANOVA found a significant difference between room temperature and cold in the liver with fructose. (B) Changes in the rate of appearance of glycerol with cold exposure assessed by stable isotope dilution (n = 9). (C) Changes in the rate of appearance of non-esterified fatty acids with cold exposure assessed by stable isotope dilution at baseline (n = 9), high fructose (n =10), and high glucose (n = 10). (D) Circulating very-low-density lipoprotein-triglycerides at room temperature and during cold (n = 10). A two-way ANOVA found a significant difference between the baseline and high-carbohydrate diets in the cold. FSF: fat signal fraction; NEFA: non-esterified fatty acids; Ra: rate of appearance; RT: room temperature; scWAT: subcutaneous white adipose tissue; VAT: visceral adipose tissue; VLDL-TG: very-low-density lipoprotein-triglycerides. Participant 13 was excluded from the analyses due to antibiotic use (potential effects on the microbiome) before the fructose and glucose diets and is shown with a square symbol shape.

We also measured circulating very low-density lipoprotein-triglycerides (VLDL-TG), which are secreted by the liver (Adiels et al., 2008). An increase in *de novo* lipogenesis due to excess glucose and, especially, fructose has been linked to increased liver fat content and circulating VLDL-TG (Alves-Bezerra and Cohen, 2018; Ter Horst and Serlie, 2017). We did observe a trend towards elevated VLDL-TG with the carbohydrate-supplemented diets compared to isocaloric diet, but this increase was only significant during cold exposure (Figure 5 D; fructose: p = 0.023; glucose: p = 0.013). Whether this is caused by increased production of VLDL-TG by the liver or by reduced tissue clearance cannot be determined with the current data.

The possible link between changes in whole-body lipolysis and lipogenesis in the liver and BAT metabolism requires further investigation. There was no correlation between baseline liver triglyceride content, or cold-induced changes in BAT triglyceride content, and BAT glucose uptake or oxidative metabolism. However, if there was an increase in VLDL-TG secretion by the liver, it could serve as an alternative energy substrate for BAT instead of glucose. Indeed, we have previously shown a direct competition between the uptake of triglyceride-rich lipoprotein-derived fatty acids and circulating glucose, with the former reducing BAT glucose uptake by more than 50% in the postprandial state (Blondin et al., 2017b).

## Discussion

Over the past decades, there has been a significant shift in our food supply system, that was accompanied by the addition of food additives such as fructose. These changes have paralleled the rise in obesity and its associated metabolic complications. This study aimed to examine whether 2-week overconsumption of fructose could modify the gut microbiome and the production of SCFA, especially acetate, and elicit changes in BAT metabolism. None of the diets, supplemented with either glucose or fructose for 2 weeks, had any effect on body composition, BAT composition, the gastrointestinal microbiome or faecal SCFA levels. Although fructose overconsumption did not change cold-induced BAT oxidative metabolism or the depletion of intracellular triglycerides, it significantly reduced BAT glucose uptake. Interestingly, this BAT metabolic disturbance occurred without any significant changes in tissue blood flow or systemic insulin resistance. Our findings demonstrate that impaired BAT glucose uptake is a very early marker of fructose-induced dysmetabolism occurring in young and otherwise healthy subjects.

We found BAT to be very sensitive to fructose-induced metabolic dysfunction, with BAT metabolic rate of glucose decreasing by 43% (38% decrease in fractional glucose extraction) within 2 weeks of fructose overfeeding. The exact cause of this reduction in BAT glucose uptake remains unclear. Increased caloric intake alone is unlikely to have caused this BAT metabolic abnormality as consuming isocaloric glucose did not change BAT glucose uptake. Neither the glucose nor the fructose-supplemented diet affected whole-body insulin sensitivity based on the homeostatic model assessment for insulin resistance (HOMA-IR), suggesting that BAT metabolic dysfunction precedes whole body insulin resistance during the development of fructose-induced metabolic dysfunction. More localized effects of insulin resistance or changes in plasma SCFA (e.g., acetate) were not assessed in this study.

The combined reduction of BAT glucose uptake with preserved BAT oxidative metabolism and intracellular triglyceride mobilization during cold exposure observed with fructose overfeeding in the healthy participants is reminiscent of our previous observation in cold-exposed individuals with T2D (Blondin et al., 2015a). Thus, 2-week fructose overfeeding provides another example of the uncoupling of BAT oxidative metabolism and glucose uptake (Blondin et al., 2017; Hankir et al., 2017; Olsen et al., 2017). Our results highlight once again the limitations of ^18^F-FDG PET as a marker of BAT thermogenesis.

Since we do not currently have a reliable non-invasive method of measuring active BAT volume in humans independent of ^18^F-FDG PET (Richard et al., 2020), we cannot determine if the impairment in glucose metabolism is accompanied by a reduction in BAT mass. For this reason, changes in the contribution of BAT to whole-body thermogenesis and glucose clearance cannot be assessed. However, we did not observe an effect of fructose overfeeding on whole-body energy expenditure, suggesting that, if there was a decrease in BAT activity, it was compensated by other thermogenic processes or it was too small to measure.

Finally, in parallel with the impairment of BAT glucose metabolism by fructose, we observed early signs of hepatic dysmetabolism with the carbohydrate-supplemented diets, and especially with high fructose. These early changes included increased lipid storage and VLDL-TG production, both early characteristic features in the development of fructose-induced fatty liver disease (Ter Horst and Serlie, 2017). Interestingly, these effects only emerged under the cold stress, as the high glucose or fructose diet did not affect hepatic lipid content or VLDL-TG levels at room temperature. The high adipose tissue intracellular triglyceride lipolytic rate stimulated in response to cold exposure resulting in increase circulating NEFA flux likely exacerbated the impact of increased hepatic *de novo* lipogenesis and, possibly reduced fatty acid oxidation that may have been induced by the high fructose diet (Low et al., 2018; Taskinen et al., 2017).

The possible link between liver and BAT impairment by fructose warrants further investigation. Notably, the increase in circulating VLDL-TG may be involved in a shift in BAT energy substrate consumption from glucose to lipids. However, this shift was not observed with the high glucose diet despite a similar increase in circulating VLDL-TG. Therefore, if a change of energy substrate occurred, it must have been driven by other factors than just an increase in circulating lipids, such as an early development of insulin resistance in BAT. Despite its apparent importance in fueling BAT thermogenesis in rodents (Bartelt et al., 2011; Berbée et al., 2015; Khedoe et al., 2015), the role of BAT lipid uptake to fuel thermogenesis in humans appears to be less prominent (Blondin et al., 2015a; Ouellet et al., 2012; U Din et al., 2016, 2018). More studies are needed to document the role of change in BAT lipid metabolism in the development of metabolic dysfunction in humans (Richard et al., 2020).

### Limitations of the study

The small sample size of our cohort is the main limitation of this study. The number of participants was voluntarily limited due to time constraints and the high cost of imaging studies. Recruitment was also difficult because of the demands placed on participants (e.g., following diets and undergoing three long and complex imaging visits). For these reasons, of the 16 participants recruited, only 10 participants completed the study. Therefore, if there were small effects of the diets on the microbiome or other outcomes, they could not be detected. Despite the small number of participants, we nevertheless demonstrated the reduction in BAT glucose uptake with fructose overfeeding without any detectable trend towards a reduction in BAT thermogenic activity. The exclusion of one subject who underwent antibiotic treatment during his participation in the study (participant 13) from the analyses did not change the results of this study.

Compliance with the diets is another possible limitation of this study. Participants were not required to fill out daily food diaries, and we thus relied on participants maintaining the same baseline diet throughout the study. However, some participants reported a loss of appetite due to the large quantities of added fructose or glucose or the large volume of beverage supplement consumed (1–1.5 L divided over three separate meals). This may have resulted in the consumption of smaller meals during fructose and glucose overfeeding, thus limiting the intended increase in caloric intake. Also, the supplementation with high quantities of glucose or fructose was designed to maximise potential effects on the microbiome, BAT, and whole-body metabolism during a short two-week period. This is not representative of normal dietary habits where smaller quantities (Powell et al., 2016; Vos et al., 2008) of added sugar (generally a combination of glucose and fructose) are consumed over long periods of time.

However, this study also includes several important strengths, including the crossover design that controlled for inter-individual variation in baseline characteristics and possible lifestyle differences, the use of a 2-week isocaloric control condition performed prior to the fructose/glucose conditions and the direct comparison between the overconsumption of fructose and glucose.

## Conclusion

Two-week fructose overfeeding did not elicit detectable changes in the microbiome or BAT thermogenesis, but impaired BAT glucose uptake, promoted liver fat storage and increased circulating VLDL-TG during cold exposure in healthy young men. Although the cause and clinical significance of this reduction in BAT glucose metabolism are unclear, our study reveals that BAT metabolic abnormality is a very early feature of fructose-induced metabolic disturbance in humans.

## Acknowledgements

The authors thank Caroll-Lynn Thibodeau, Maude Gérard, Éric Lavallée, Etienne Croteau, Esteban Espinosa, Jean-Philippe Pelletier, Luc Tremblay, Guillaume Gilbert, Christophe Noll, Lucie Bouffard, Zaccary Corradini-Carriere, and all the MR technologists involved in this study for their excellent technical assistance. They also thank the staff of the Farncombe Metagenomic Facility (Hamilton, CA) for support in library sequencing. They also thank Réjean Lebel for his help revising this manuscript. This work was supported by the Canadian Institutes of Health Research (grant number 144625-1, 299962 and PJT 159758). G.R. is a recipient of a NSERC Alexander-Graham-Bell Ph.D. scholarship. D.P.B. holds the GSK Chair in Diabetes of Université de Sherbrooke and a Fonds de Recherche du Québec-Santé (FRQS) J1 salary award. S.A.S. is a recipient of a CIHR Vanier Ph.D. scholarship. A.C.C. holds the Canada Research Chair in Molecular Imaging of Diabetes.

## Author contributions

Conceptualization, A.C.C., M.L., D.P.B., J.D.S., G.R.S., K.M.M., E.E.T, and B.G.; Methodology: A.C.C., M.L., D.P.B., G.R., J.D.S., G.R.S., K.M.M., M.G.S., J.D.S., E.E.T, M.F., S.P., F.F., S.D., and B.G.; Investigation: A.C.C., D.P.B., G.R., S.A.S., L.R., M.E.F., M.F., S.P., F.F., E.E.T., and B.G.; Writing – first draft: G.R., D.P.B., A.C.C., S.A.S.; Writing – review & editing: S.A.S., G.R.S., K.M.M., M.G.S., J.D.S., A.C.C., B.G., M.L., S.D., M.F.; Visualization: G.R., S.A.S.; Funding Acquisition: A.C.C., J.D.S., G.R.S., and K.M.M.

## STAR METHODS

### KEY RESOURCES TABLE

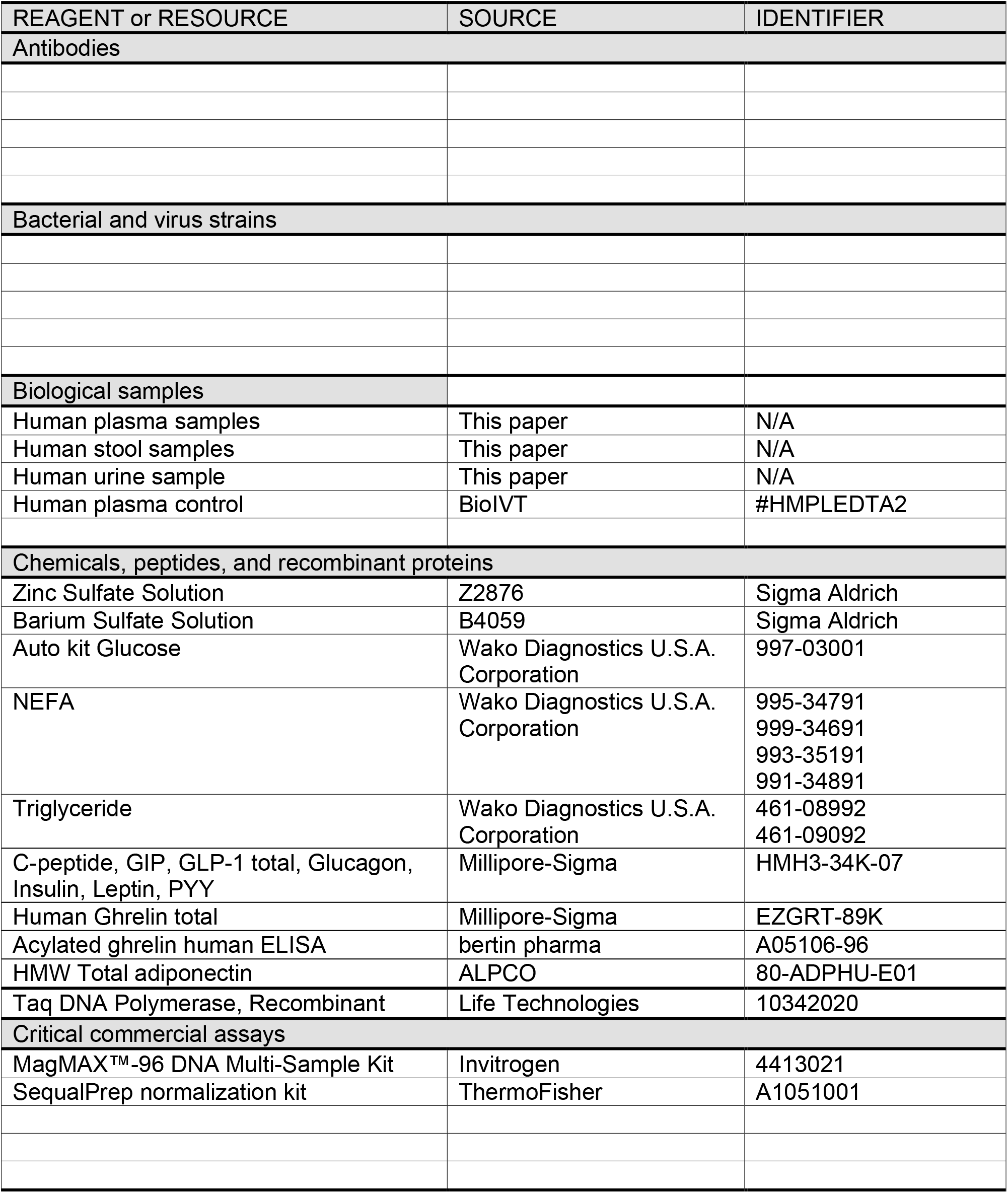
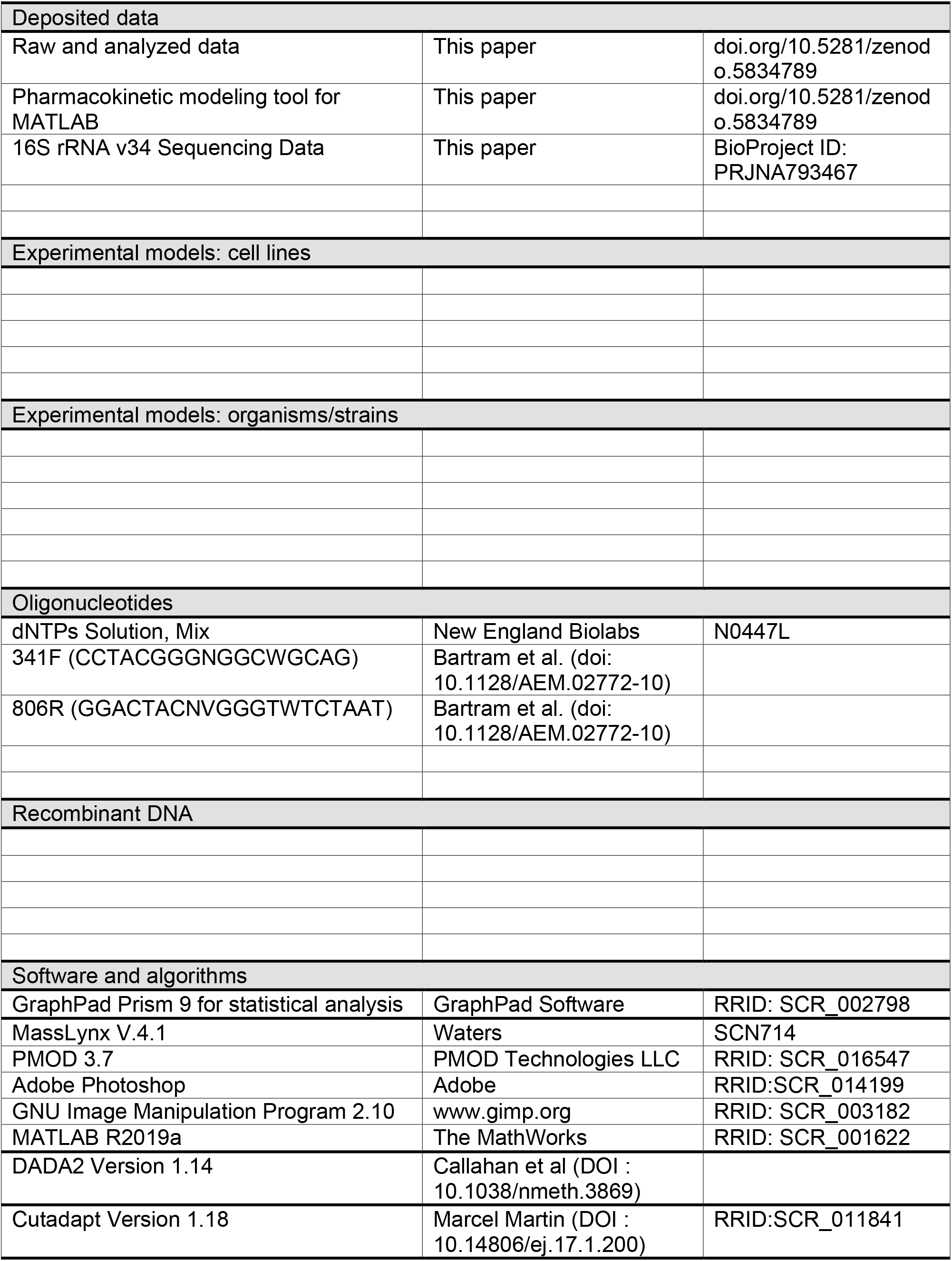
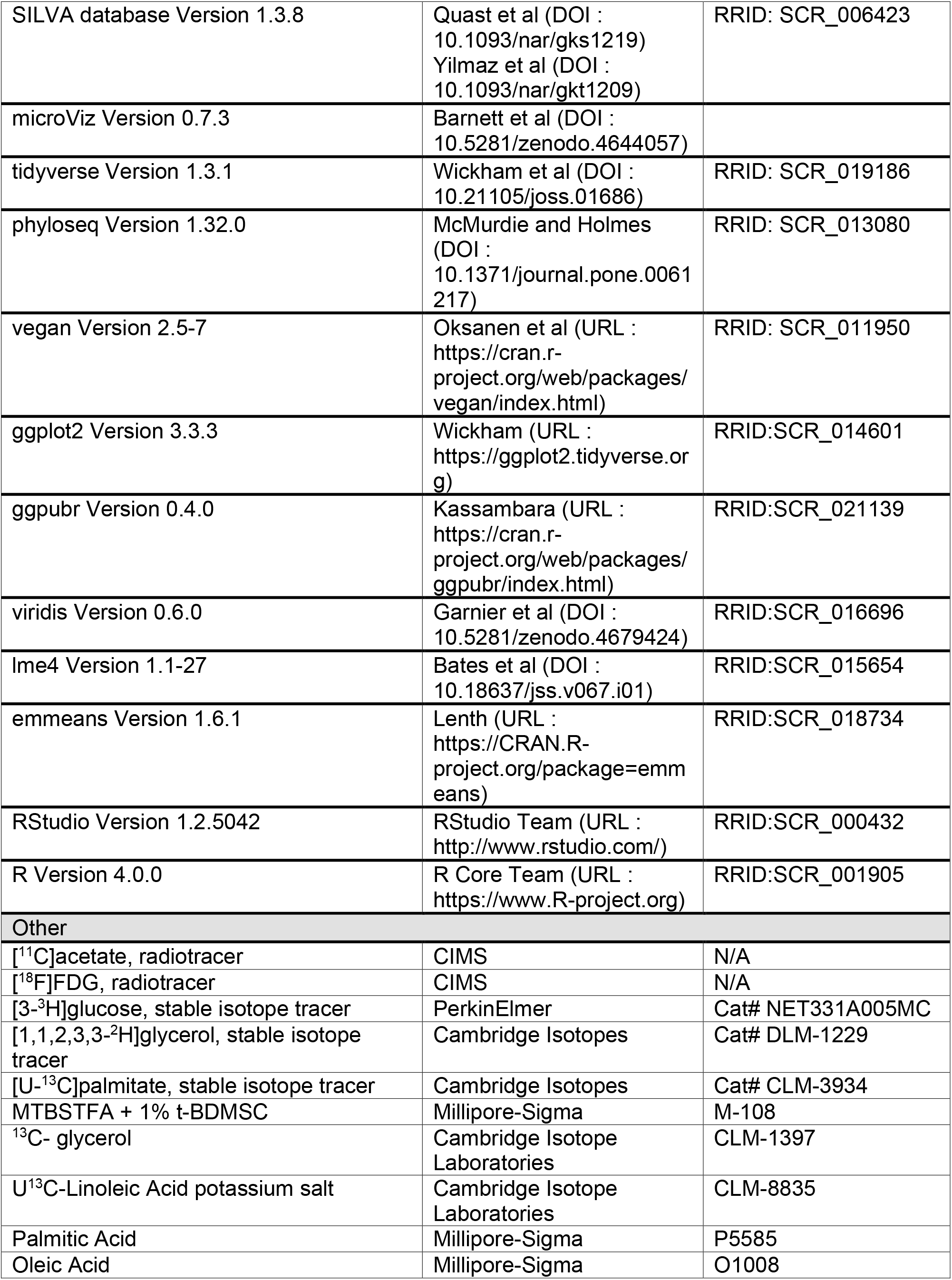
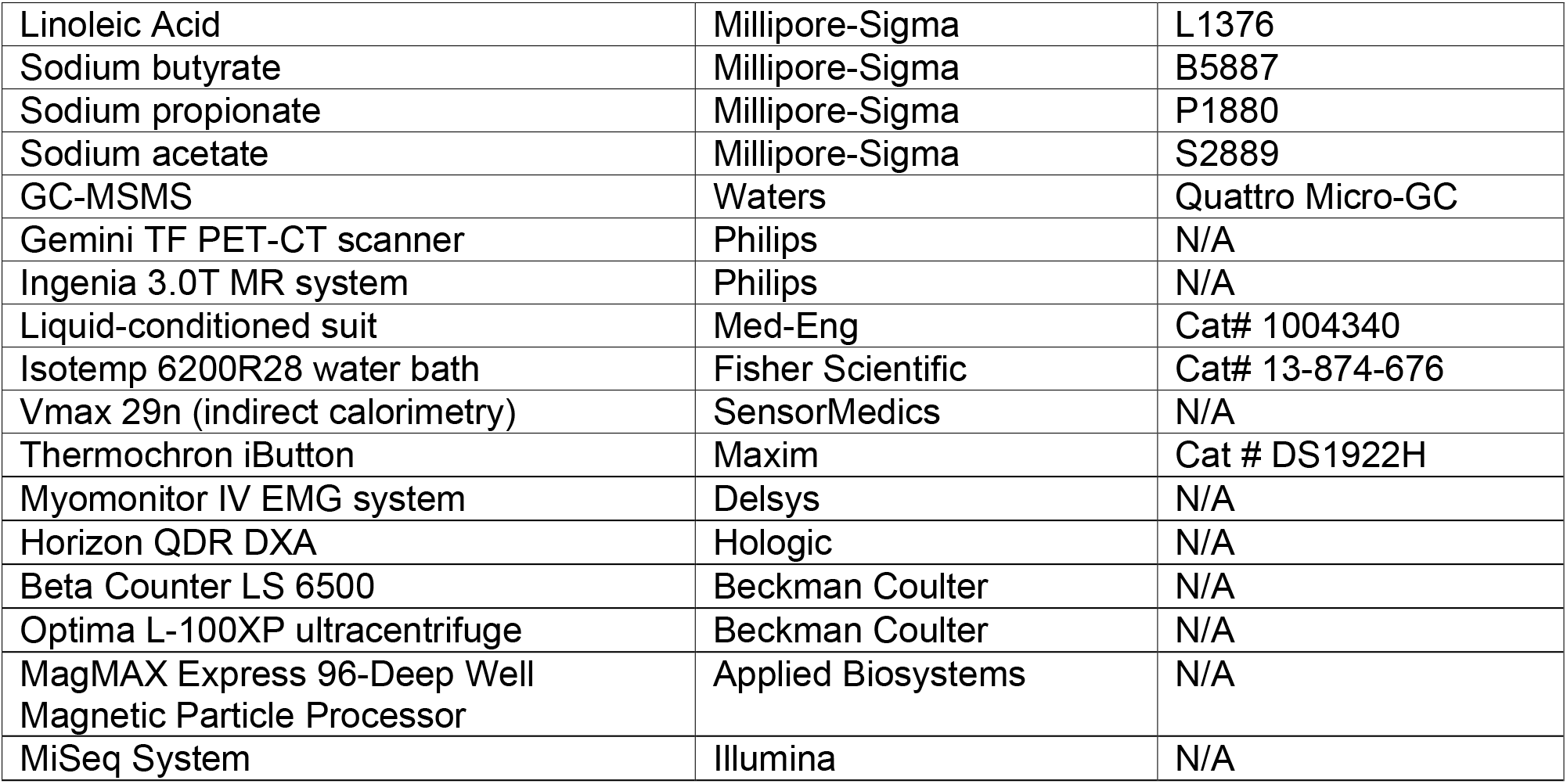

## RESOURCE AVAILABILITY

### Lead contact

Further information and requests for resources and reagents should be directed to and will be fulfilled by the lead contact, André Carpentier.

### Materials availability

This study did not generate unique reagents.

### Data and code availability

The processed data generated during this study as well as the MATLAB code used to perform the pharmacokinetic modeling of ^18^F-FDG and ^11^C-acetate are available at doi.org/10.5281/zenodo.5834789. Anonymized image files are voluminous and, therefore, not deposited in any public repository, but can be made available on reasonable request to the lead contact.

## EXPERIMENTAL MODEL AND SUBJECT DETAILS

### Human Subjects

The protocol received approval from the Human Ethics Committee of the Centre de Recherche du Centre hospitalier universitaire de Sherbrooke. Written informed consent was obtained from all participants in accordance with the Declaration of Helsinki. The study was registered on ClinicalTrials.gov (NCT03188835). A total of 16 male participants were recruited for this study (Supplemental Figure S1). However, 5 withdrew before undertaking any experimental procedures. Of the remaining 11 participants, 1 withdrew following the baseline imaging protocol and 1 was excluded from analyses of carbohydrate-enriched diets (but not baseline) due to antibiotic use potentially affecting the microbiome. Metabolic data for this last participant were still analyzed and shown on graphs using a different symbol shape. Characteristics of participants can be found in Table S1. The inclusion criteria were as follows: (i) male; (ii) age between 20 and 40 years-old; (iii) BMI < 30 kg/m^2^; (iv) normal glucose tolerance (blood glucose < 7.8 mmol/L 2 h post 75 g oral glucose tolerance test); (v) no first-degree familial history of type 2 diabetes (parent or sibling). Exclusion criteria comprised: (i) fasting plasma triglycerides > 5.0 mmol/L or history of total cholesterol > 7 mmol/L; (ii) consuming more than 2 alcoholic beverages per day; (iii) smoking more than 1 cigarette per day; (iv) history of total cholesterol level > 7 mmol/L, of cardiovascular disease or hypertensive crisis; (v) treatment with fibrates, thiazolidinedione, insulin, beta-blocker or other drugs with effects on insulin resistance or lipid metabolism (exception for anti-hypertensive drugs, statins or metformin); (vi) presence of a non-controlled thyroid disease, renal or hepatic disease, history of pancreatitis, bleeding disorder, cardiovascular disease or any other serious medical condition; (vii) history of serious gastrointestinal disorders (malabsorption, peptic ulcer, gastroesophageal reflux having required a surgery, etc.); (viii) contraindication to MR imaging (e.g., metal implant) or PET/CT imaging (e.g., having undergone a scan in the last 12 months); (ix) chronic administration of any other medication.

## METHOD DETAILS

### Study design

Supplemental Figure S2 summarizes the study protocols. Briefly, participants were asked to follow three 14-day dietary protocols. The first protocol (A) consisted of a balanced diet for baseline measures. The other two protocols consisted of the same baseline diet supplemented with a high fructose (B) or glucose (C) drink, each representing 25% of energy intake. Protocols B and C were randomized and separated by a 1-month washout period during which participants followed the baseline diet. After each diet protocol, participants provided stool samples for microbiome analysis and underwent a metabolic study day with cold exposure to stimulate BAT thermogenesis. During the metabolic study days, body composition was assessed by dual-energy X-ray absorptiometry (DXA) and magnetic resonance (MR). Whole-body energy expenditure and substrate metabolism were measured by indirect calorimetry and stable isotope tracers. Tissue-specific glucose uptake was assessed by ^18^F-FDG PET, and oxidative metabolism was assessed by ^11^C-acetate PET. Participants were fasted for 12 h and abstained from strenuous physical activity for 48 h prior to metabolic study days.

#### Measures of body composition and energy expenditure

Body composition was assessed by DXA (Hologic) and bioelectrical impedance (Tanita). Whole-body energy expenditure at rest was determined by indirect respiratory calorimetry with correction for protein oxidation (Haman et al., 2002) for a duration of 20 minutes per measure. These measures were taken on visits 2, 3, 4, 5 and 6. On imaging visits (visits 3, 4 and 6), the indirect calorimetry (SensorMedics Vmax) was performed once at room temperature (before cooling) and twice during cold exposure (the reported cold exposure data is the average of both measures). The molar rate of fatty acid oxidation was determined by converting the rate of triglyceride oxidation (mg/min) to its molar equivalent (μmol/min), assuming the average molecular weight of human triglyceride is 860 g/mol and multiplying the molar rate of triglyceride oxidation by three, as each mole of triglyceride contains three moles of fatty acids (Wolfe and Chinkes, 2005). Energy potentials of 16.3, 40.8 and 19.7 kJ/g were used to calculate the amount of heat produced from glucose, lipid, and protein oxidation, respectively, and converted to a rate in kJ·min^−1^. The level of physical activity of each participant was assessed with a uniaxial accelerometer (Caltrac) on 3 consecutive days during each diet protocol (before visits 3, 4 and 6).

#### Diets

The baseline, isocaloric, meal plans were designed by a registered dietician (S.D.) following the recommendations of the 2007 Canada’s Food Guide (~50% carbohydrates, ~30% lipids and ~20% proteins) as well as the dietary preferences and restrictions of each participant. Daily caloric needs were estimated based on resting metabolism (indirect calorimetry) and physical activity (uniaxial accelerometer). Alcohol and dietary supplements likely to affect the gastrointestinal microbiome were prohibited. Participants were asked to refrain from changing their dietary or exercise habits during the study. However, they were not asked to fill out food diaries or otherwise supervised to ensure compliance.

The fructose (protocol B) or glucose (protocol C) supplements were provided in powder form (Sigma-Aldrich) that participants dissolved in 1 to 1.5 L of water and drank throughout the day. Addition of pure lemon or orange juice (no artificial sweeteners) was suggested to make the drink more palatable. To improve tolerance, the amount of glucose or fructose consumed was gradually increased during the first 3 days of the study (15%, 20% and 25% of estimated daily energy requirements) and remained stable for the remaining 11 days (174 ± 37 g/day, equivalent to 655 ± 138 kcal/day).

Finally, participants fasted (water allowed) from 20 h 00 on the evening before the metabolic study until the end of procedures at approximately 16 h 30 the following day.

#### Stool sample collection and analysis

Stool samples were provided by participants after each diet protocol. Samples were kept frozen at −80 °C until processed. Before analysis, samples were thawed either at 4 °C (faecal short-chain fatty acid content) or room temperature (microbiome composition), and remaining portions were refrozen. Stool samples underwent analysis for SCFA (acetate, propionate, and butyrate) at Université de Sherbrooke, as well as microbiome composition at McMaster University.

Faecal SCFA content was measured by gas chromatography-tandem mass spectrometry (GC-MS/MS, Agilent GC model 7890A and Waters Quattro Micro) based on methods developed for blood samples (Pouteau et al., 2001; Tsukahara et al., 2014). The column was an Agilent DB-5ms Ultra Inert (30 m, 0.25 mm, 0.25 μm) with helium as the carrier gas. Each sample was analyzed in triplicate. All samples from a same participant were analyzed at the same time to limit variability. Approximately 150 mg of sample was transferred to a 2 mL polypropylene microtube with glass beads and weighted to obtain wet weight. Then, 1 mL of 0.2% HCL was added, and the sample was shaken on ice for 20 minutes before centrifugation (9400 g, 5 minutes). The supernatant was transferred to a 2 mL polypropylene tube and recentrifuged (9400 g, 2 minutes). The pellet was conserved for estimation of dry weight. For sample analysis, 20 μL of diluted internal standard (acetate-d3 25 mM, propionate-d5 5 mM or butyrate-d5 5 mM) was added to a 2 mL crimp top vial, followed by 250 μL of 0.4% HCL, 180 μL of deionised water, 50 μL of the supernatant, and 1 mL of dichloromethane. The tube was closed with the crimp cap and mixed with a shaker (10 minutes, room temperature) before being centrifuged upside down (800 g, 5 min). Approximately 600 μL of the dichloromethane phase was drawn from the upside-down vial with a glass syringe (avoiding any water). The dichloromethane solution was transferred to a 2 mL glass vial containing 400 mg of dry sodium sulfate. The glass vial was closed and allowed to sit for one hour at room temperature. Finally, 20 μL of the derivatization reagent (MTBSTFA + 1% t-BDMCS reagent diluted 1:100 in dichloromethane) was added to a 2 mL screw-tread vial with a 200 μl, 6 x 29 mm glass insert (mandrel precision point, 33 glass, with plastic spring; Chromatographic Specialties), followed by 200 μL of sample extract (dichloromethane phase). The sample was then injected (3 μL) in the GC-MS/MS in split mode and using positive electron impact ionization for mass detection.

Genomic DNA was extracted as described previously (Stearns et al., 2015) with some modifications (Szamosi et al., 2020). Samples were transferred to screw cap tubes containing 2.8 mm ceramic beads, 0.1 mm glass beads, GES and sodium phosphate buffer. Samples were bead beat and centrifuged as described, and the supernatant was further processed using the MagMAX Express 96-Deep Well Magnetic Particle Processor from Applied Biosystems with the multi-sample kit (Life Technologies #4413022).

Purified DNA was used to amplify the v34 region of the 16S rRNA gene by PCR. 50 ng of DNA was used as template with 1U of Taq Polymerase, 1x Taq buffer, 1.5 mM MgCl2, 0.4 mg/mL BSA, 0.2 mM dNTPs, and 5 pmoles each of 341F (CCTACGGGNGGCWGCAG) and 806R (GGACTACNVGGGTWTCTAAT) Illumina adapted primers, as described by Bartram and colleagues (Bartram et al., 2011). The reaction was carried out at 94 °C for 5 minutes, 5 cycles of 94 °C for 30 seconds, 47 °C for 30 seconds, and 72 °C for 40 seconds, followed by 25 cycles of 94 °C for 30 seconds, 50 °C for 30 seconds and 72 °C for 40 seconds, with a final extension of 72 °C for 10 minutes. Resulting PCR products were visualized on a 1.5% agarose gel. Positive amplicons were normalized using the SequalPrep normalization kit (ThermoFisher #A1051001) and sequenced on the Illumina MiSeq platform at the McMaster genomics facility.

Reads were processed using DADA2 (Callahan et al., 2016). First, Cutadapt was used to filter and trim adapter sequences and PCR primers from the raw reads with a minimum quality score of 30 and a minimum read length of 100 bp (Martin, 2011). Sequence variants were then resolved from the trimmed raw reads using DADA2, an accurate sample inference pipeline from 16S amplicon data. DNA sequence reads were filtered and trimmed based on the quality of the reads for each Illumina run separately, error rates were learned, and sequence variants were determined by DADA2. Sequence variant tables were merged to combine all information from separate Illumina runs. Bimeras were removed and taxonomy was assigned using the SILVA database version 1.3.8.

We filtered to all ASVs assigned to the kingdom bacteria and removed all ASVs assigned to the kingdom eukaryota, the order chloroplast and the family mitochondria. After filtering, 1,720,471 total reads (average: 61,445; range: 11,449–124,952) with 2517 total ASVs (average 199 ASVs/sample; range: 61–341) remained.

Please refer to the *Quantification and statistical analysis* section for details on the analysis methods.

#### BAT stimulation by cold exposure

On metabolic study days, subjects underwent a 3-hour cooling protocol (Blondin et al., 2017). After the morning MRI session, subjects, wearing only shorts, were instrumented with autonomous wireless temperature sensors (Thermochron iButton model DS1922H, Maxim) placed on 12 sites to measure mean skin temperature (Hardy et al., 1938): forehead, upper back, lower back, abdomen, forearm, quadricep, hamstring, gastrocnemius, anterior tibialis, chest, bicep, and hand. Surface electromyography electrodes (Delsys, EMG System) were placed on the belly of eight muscles known to contribute significantly to shivering during cold exposure (Bell et al., 1992; Blondin et al., 2015b; Haman et al., 2004a, 2005): *m.* trapezius superior, *m.* sternocleidomastoid, *m.* pectoralis major, *m.* deltoideus, *m.* biceps brachii, *m.* rectus abdominis, *m.* vastus lateralis, *m* rectus femoris, and *m.* vastus medialis. To normalize the shivering measures, subjects performed a series of isometric maximal voluntary contractions for each muscle. Finally, participants were asked to put on a liquid conditioned suit (Three Piece, Med-Eng).

Cooling was started ~12 h 00 and maintained for 180 min. During this period, water at 18 °C was perfused through tubing in the suit with a temperature- and flow-controlled circulation bath (Isotemp 6200R28, Fisher Scientific). Shivering and skin temperature were monitored continuously throughout the experiment. Shivering intensity and shivering pattern were determined as previously described (Blondin et al., 2015b; Haman et al., 2004a, 2004b, 2005). Mean skin temperature was calculated using an area-weighted mean from the wireless temperature sensors.

#### Energy substrate turnover

On metabolic study days, after the morning MRI, participants were asked to empty their bladders and indwelling catheters (18G) were placed in the antecubital vein of both arms for blood sampling (right arm) and tracer infusion (left arm). A primed (3.3 × 10^6^ dpm) continuous infusion (0.56 × 10^6^ dpm min^−1^) of [3-^3^H]-glucose was initiated 150 min before the start of cold exposure to determine the plasma glucose appearance rate (Ra_glucose_) (Ouellet et al., 2012; Vella and Rizza, 2009). A bolus of [1-^13^C]-NaHCO3 (1.2 μmol·kg^−1^), followed by continuous infusion of [U-^13^C]-palmitate (0.01 μmol·kg^−1^·min^−1^ in 100 mL 25% human serum albumin), and a primed (1.6 μmol·kg^−1^) continuous infusion (0.1 μmol·kg^−1^·min^−1^) of [1,1,2,3,3-^2^H]-glycerol were started 60 min before the start of cold exposure to determine plasma NEFA appearance rate (Ra_NEFA_) and plasma glycerol appearance rate (Ra_glycerol_).

RaNEFA was calculated by multiplying the plasma palmitate appearance rate by the fractional contribution of palmitate to total plasma NEFA concentrations. Blood samples were collected for dosing of tracer enrichment and other metabolites before starting the infusion, before cooling, and every hour thereafter. Samples of expired air were also taken every hour to measure the enrichment in ^13^CO_2_. Urine was collected at the end of the baseline period, 75 min into cold exposure and at the end of cold exposure to measure urea production. Rates of appearance for glucose, non-esterified fatty acids, and glycerol were reported for room temperature (t = 0, just before cold exposure) and the steady state of the cooling period (t = 120 and 180 min).

#### Imaging of triglyceride content by magnetic resonance imaging

On metabolic study days, MRI and MRS were performed in the morning (before cooling, at approximately 7 h 30) and evening (after cooling, at approximately 16 h 00) on a 3T Philips Ingenia scanner (Philips Healthcare). On both occasions, triglyceride content was assessed in the supraclavicular region and in the abdominal region.

Participants, wearing only the standard hospital gown, were placed on a dedicated table (Philips Healthcare) compatible with the MR and PET scanners. They were immobilized with a vacuum cushion (Vac-Lok; Civco Radiotherapy) to ensure consistent positioning throughout the day. In the morning, participants were wrapped in a warm blanket to ensure participants were not cold.

Images were acquired with an 8-channel torso surface coil (dStream; Philips) covering the neck, torso, and abdomen. Tables S2 and S3 summarize the MRI and MRS sequence parameters. Briefly, an anatomical T1-weighted image of the supraclavicular region was taken to localize BAT. Three 8 mm^3^ voxels were placed in the supraclavicular fat for BAT proton spectroscopy. Voxel placement aimed at avoiding blood vessels and surrounding muscles while covering the largest surface possible (Supplemental Figure S3). Since supraclavicular fat depots are bilateral, at least one voxel was placed on each side. To quantify fat content, mDixon Quant images were acquired covering the neck and shoulders. Similar sequences were performed for the liver region, except that a single 25 mm^3^ voxel was placed in the right lobe of the liver, and a single 10 mm^3^ voxel was placed in the subcutaneous fat of the lower back. Liver spectroscopy, Dixon imaging, and T1-weighted image of the liver were performed in breath hold (inspiration), other sequences were performed with free breathing.

#### Imaging of glucose metabolism and oxidative metabolism by PET/CT

A dynamic ^11^C-acetate PET scan (Philips Gemini TF) was performed at room temperature at approximately 11 h 30 only for protocol A (baseline). For all protocols, a dynamic ^11^C-acetate scan was performed at approximately 13 h 30 after 90 minutes of cooling. Each scan started with the injection of ^11^C-acetate (~185 MBq, over 20 s with a 20 s saline flush) and was performed as a 30-minute list-mode acquisition. A dynamic ^18^F-FDG scan was performed at approximately 14 h 00 with a 2-minute background, injection of the radiotracer (~125 MBq, over 20 s with a 20 s normal saline flush) and 30-minute list-mode acquisition. Finally, a whole-body (head to knee) static PET scan was performed at approximately 15 h 00, after the 3-hour cooling period. This static scan measured ^18^F-FDG distribution in different tissues without additional radiotracer injection.

Low-dose CT scans were obtained for anatomical reference and attenuation correction. A CT of the cervicothoracic region (120 kVp, 49 mA, 30 mAs) was done before each dynamic ^11^C-acetate scan, and a whole-body CT (120 kVp, 30 mA, 16 mAs) was done before the whole-body ^18^F-FDG scan. CT acquisition was not needed for the dynamic ^18^F-FDG scan because the participant was in the same position as for the previous ^11^C-acetate scan.

#### Image processing

MR images (Supplemental Figure S4) were reconstructed online. Dynamic PET images were reconstructed with a 3D RAMLA algorithm. Time frames for ^11^C-acetate were: 24 × 10 s, 12 × 30 s, and 4 × 300 s. Time frames for ^18^F-FDG were: 1 x 2 min (background), 12 × 10 s, 10 × 45 s, 7 × 90 s, and 2 ×300 s. The static ^18^F-FDG image was reconstructed with a BLOB-OS-TF algorithm. CT reconstruction was performed with filtered back projection. Standard corrections for decay, attenuation (CT-based), scattering (single scatter simulation) and random coincidences (delayed window) were performed for PET. PET data was expressed in units of standardized uptake value 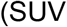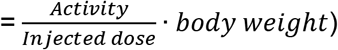.

Regions of interest were segmented manually in PMOD (v. 3.7, Bruker). Axial CT images were used as anatomical references for PET segmentation. Axial T1-weighted images were used as anatomical reference for MRI segmentation. PET and MR images were also registered and segmented together, but this method proved less reliable due to small registration errors, and results were not presented in this article. The BAT region included all fat visible in the neck and supraclavicular fossa on both sides. Other BAT depots (e.g., paraspinal) were not segmented because they are not visible for all participants. Other segmented tissues include muscles (*m.* sternocleidomastoideus, *m.* trapezius, *m.* deltoideus, and *m.* pectoralis major), subcutaneous white adipose tissue (lower back for MRI and cervical for PET), abdominal visceral adipose tissue (excluding perirenal adipose tissue), and liver. For pharmacokinetic modeling, the arterial input function was extracted from a spherical region (diameter: 10 mm) in the aortic arch.

For static images, such as fat signal fraction maps from Dixon MRI, CT radiodensities, and static ^18^F-FDG PET, average values were calculated for each region of interest. For dynamic PET imaging, time-activity curves were extracted for each region of interest. The blood signal for ^11^C-acetate was then corrected to exclude the contribution of metabolites (Buck et al., 1991).

Pharmacokinetic modeling was performed in MALTB (The MathWorks, R2019a). A four-compartment, two-tissue, model (Richard et al., 2019) was applied to the ^11^C-acetate signal to derive the rates of uptake (*K*_1_ [ml·g^−1^·min^−1^]), oxidation (*k*_2_ [min^−1^]) and lipid synthesis (*k*_3_ [min^−1^]). For ^11^C-acetate under cold condition, the blood volume fraction was constrained based on data from the ^18^F-FDG three-compartment model (Richard et al., 2021). Fractional glucose uptake (*K*_i_ [min^−1^]) was also obtained from Patlak graphical analysis (Patlak and Blasberg, 1983) and converted to metabolic rate of glucose using the plasma glucose level (MRGlu = *K*_i_·glycemia·LC^−1^·100 [μmol·100 g^−1^·min^−1^]). This conversion assumes a lump constant (LC) value for ^18^F-FDG compared with endogenous plasma glucose which was set at 1.14 for fat and 1.16 for muscles (Peltoniemi et al., 2000; Virtanen et al., 2001).

Finally, MR spectra were processed in jMRUI and quantified with AMARES (Vanhamme et al., 1997). For BAT and scWAT, 6 gaussian peaks were considered: 0.9 ppm methyl, 1.3 ppm methylene/β-carboxyl, 2.1 ppm α-olefin/α-carboxyl, 4.2 ppm glycerol, 4.7 ppm water, and 5.3 ppm olefin/glycerol. For the liver, only the water and methylene peaks were visible. Correction for T1 and T2 decay was performed for each resonance. For T1, data from the literature was used (Hamilton et al., 2011; Peterson et al., 2021). For T2, correction was based on the multi-echo data for the liver and on the literature for BAT and scWAT (Hamilton et al., 2011; Peterson et al., 2021). Voxels with poor shimming or excessive signal contamination by other tissues were rejected. Values obtained for the different BAT voxels in a same participant and condition were averaged. The number of double bonds and methylene-interrupted double bonds were calculated for BAT and scWAT based on the method by Hamilton and colleagues (Hamilton et al., 2011).

#### Biological Assays

Individual plasma NEFA (palmitate, linoleate, oleate) [U-^13^C]-palmitate enrichment and [1,1,2,3,3-^2^H]-glycerol enrichment were measured by GC-MS/MS (Agilent GC model 7890A and Waters Quattro Micro). The column was an Agilent DB-5ms Ultra Inert (30 m, 0.25 mm, 0.25 μm) with helium as the carrier gas. The precipitation method was based on that of Patterson and colleagues (Patterson et al., 1999) with some modifications. An aliquot of plasma (40 μL) was mixed with internal standard of ^13^C_18_-linoeate (20 μL of 50 μM in acetone) and ^13^C-glycerol (20 μL of 500 μM in water). Then, acetone (500 μL) was added, and the mixture was cooled at −20 °C to allow protein precipitation. After centrifugation, (3500 g, 10 minutes), the supernatant was extracted, then mixed with deionised water (500 μL) and hexane (500 μL). After another centrifugation (3500 g, 10 minutes), the two phases were separated, and dried under vacuum. The organic phase residue, which contains free fatty acids was derivatized with a solution of MTBSTFA + 1% t-BDMCS (8.2 mg, 34 μmol) and imidazole (54 ng, 0.8 nmol) in DMF. Aqueous phase residue for glycerol quantification was derivatized with a solution of MTBSTFA + 1% t-BDMCS (20.7 mg, 86 μmol) and imidazole (136 ng, 2 nmol) in DMF (40 μL). After incubation (80 °C, 1 hour), samples were cooled down and then injected (1 μL) in the GC-MS/MS in split mode and using positive electron impact ionization for mass detection.

Tritium from [1-^3^H]-glucose enrichment was detected by liquid scintillation spectrometry (Beckman Coulter LS 6500).

Plasma glucose, total NEFA triglycerides, cortisol, TSH, free T3 and free T4 were measured using specific radioimmunoassays and colorimetric assays. Plasma C-peptide, GIP, total GLP-1, glucagon, insulin, and leptin were measured using Luminex xMAP-based immunoassays (Millipore).

## QUANTIFICATION AND STATISTICAL ANALYSIS

### Statistical Analysis

All microbiome analyses were conducted in RStudio (version 1.2.5042; April 2020) with R (version 4.0.0; April 2020) using the microViz, tidyverse, phyloseq, vegan, lme4, and emmeans packages for analysis, and ggplot2, ggpubr, and viridis for visualization. For α-diversity calculations, samples were rarefied to 11,449 sequence reads per sample. Differences in α-diversity between our metabolic measures of interest were assessed using generalized mixed-effects model analysis with participants used as the random effect variable and diet and sequence used as fixed effects. To measure β-diversity, samples were normalized by centred log-ratio transformation. Differences in β-diversity were assessed using permutational multivariate analysis of variance (PERMANOVA) assessing for differences by diet.

Metabolic data (including imaging data, blood chemistry, indirect calorimetry, and stable isotopes) are expressed as mean with 95% confidence interval or median with interquartile range. Analyses were performed in Prism (GraphPad; 9.0.2). Normality of data was assessed by the Shapiro-Wilk test. For normally distributed data, the effects of diets and cold on whole-body energy expenditure and substrate turnover as well as specific tissue composition and metabolism were assessed by a two-way ANOVA with Bonferroni correction for multiple comparisons. Otherwise, for data that was not normally distributed, the Friedman test was used with Dunn’s correction for multiple comparisons. Correlation and linear regression analyses were applied to investigate the relationships between BAT metabolic parameters and other metabolic and hormonal cofactors. The significance threshold was set at p < 0.05.

## ADDITIONAL RESOURCES

ClinicalTrials.gov identifier NCT03188835

