## Supplemental figures and tables for "High-fructose feeding suppresses cold-stimulated brown adipose tissue glucose uptake in young men independently of changes in thermogenesis and the gut microbiome"

### SUPPLEMENTAL INFORMATION

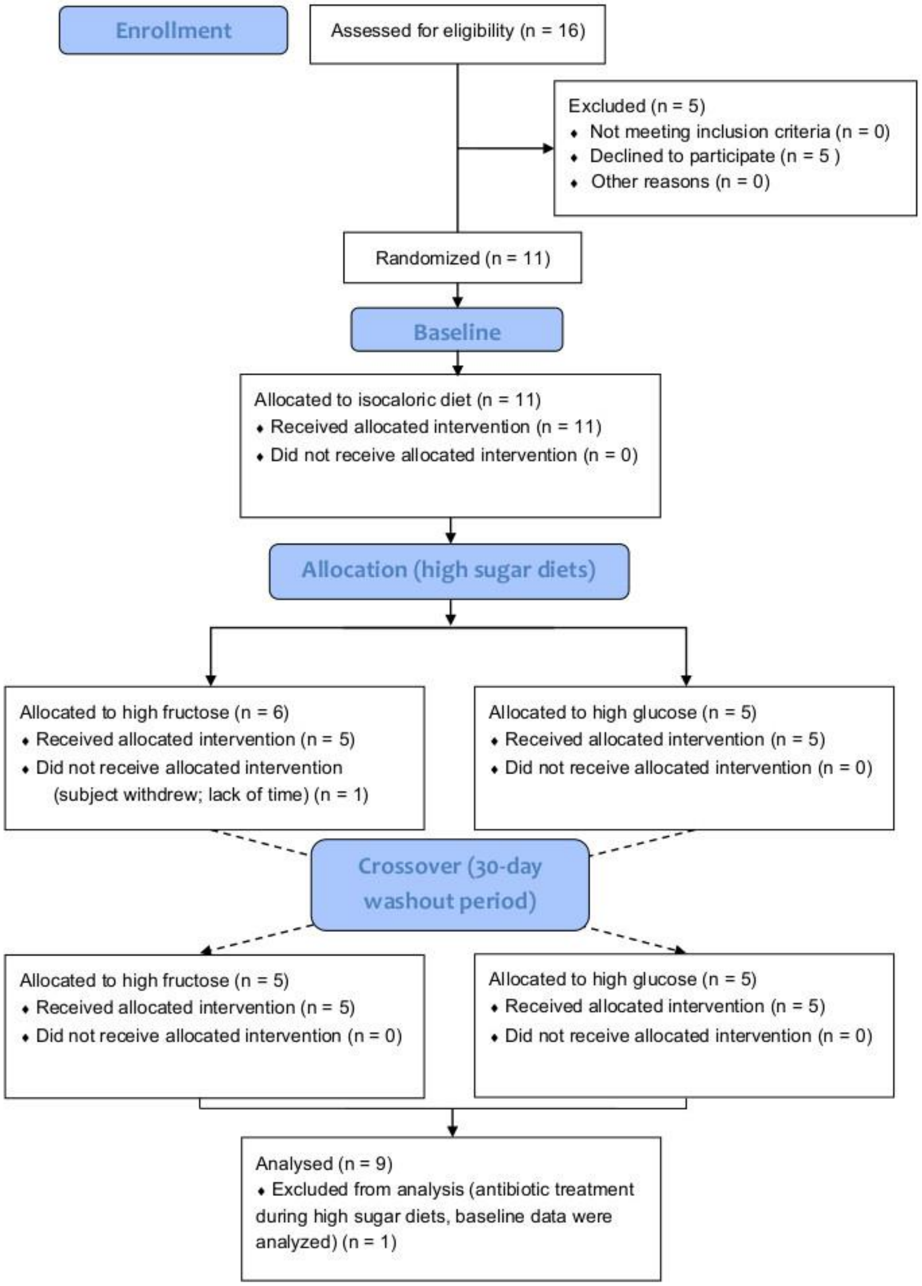


**Figure S1.** CONSORT flow diagram illustrating participant enrollment, allocation, and analysis for the study.

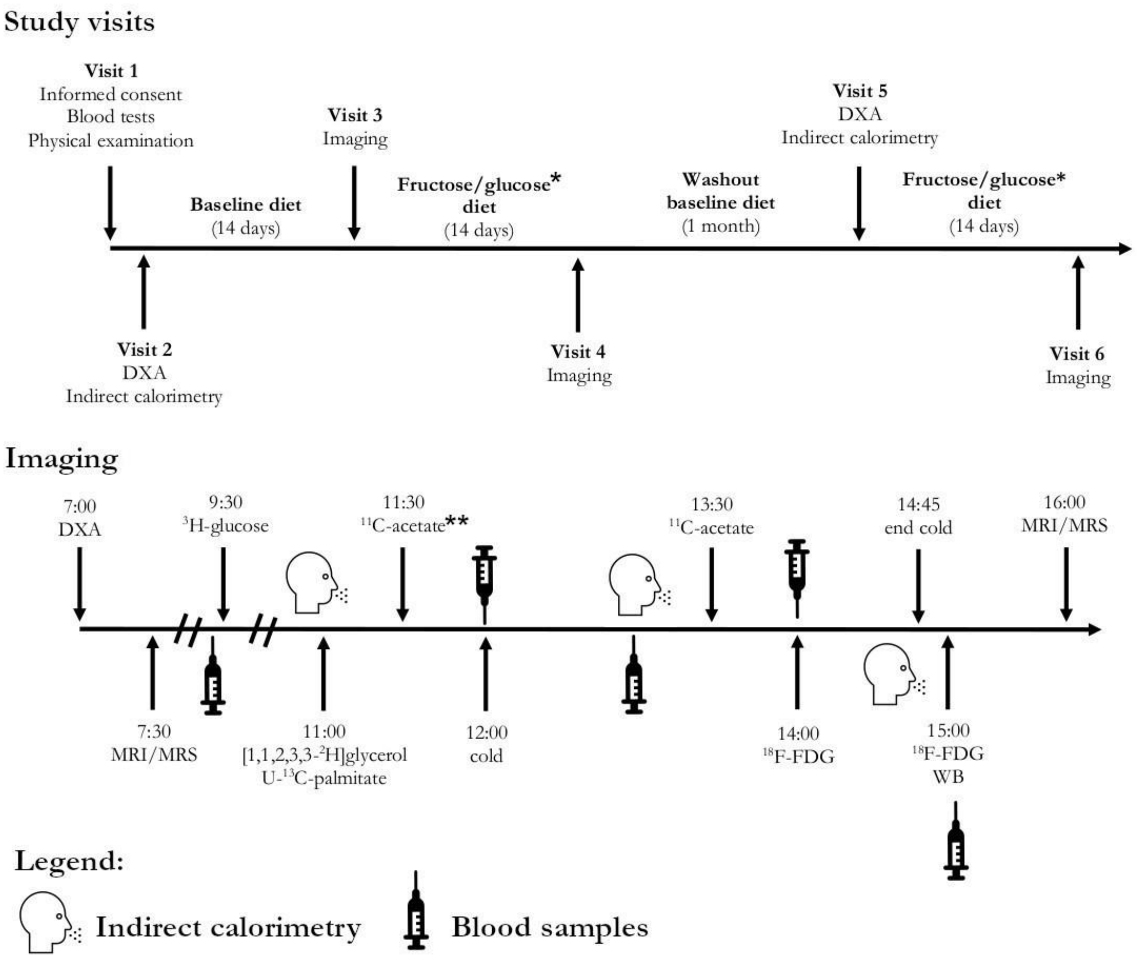


**Figure S2.** Summary of study protocol. The study had 6 visits including three imaging studies. Body composition was assessed before and after each diet. Stool samples were collected after each diet, before the imaging study. *Fructose and glucose diets were randomized. **The 11:30 (room temperature) $^{11}$C-acetate scan was only performed at baseline. DXA: dual-energy X-ray absorptiometry; FDG: fluorodeoxyglucose; MRI: magnetic resonance imaging; MRS: magnetic resonance spectroscopy; WB: whole-body.

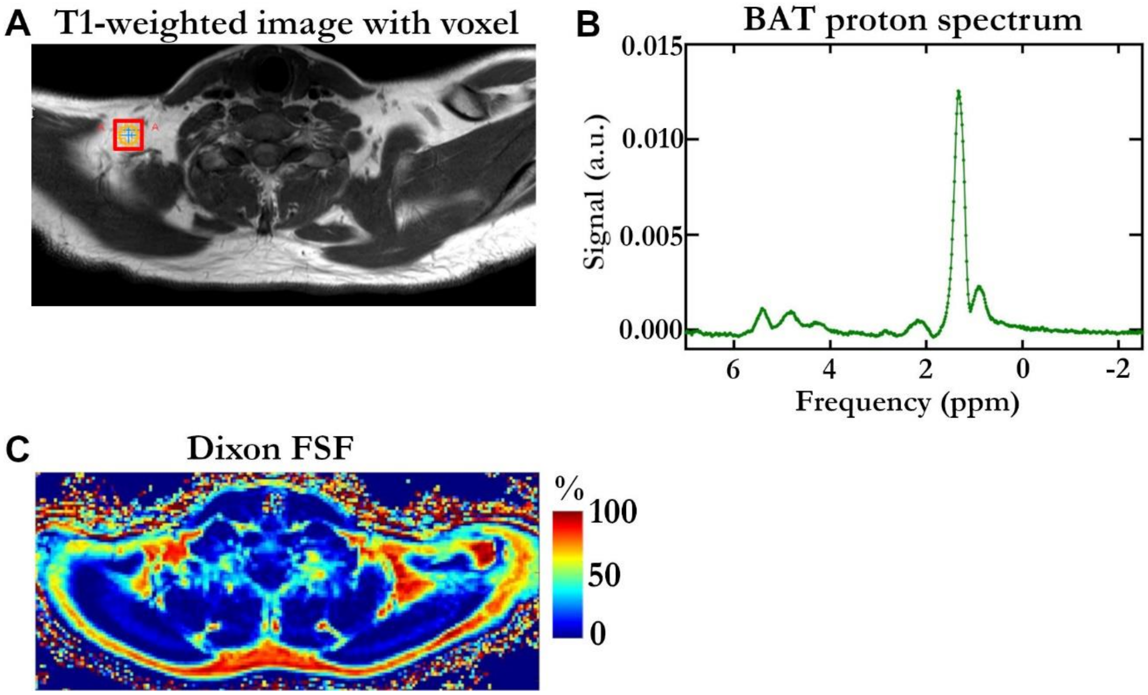


**Figure S3.** Typical magnetic resonance images and spectrum. (A) Anatomical T1-weighted image with spectroscopy voxel placement in brown adipose tissue. (B) Typical brown adipose tissue proton spectrum. (C) Fat signal fraction map from mDixon. See Tables S2 and S3 for sequence parameters. BAT: brown adipose tissue; FSF: fat signal fraction.

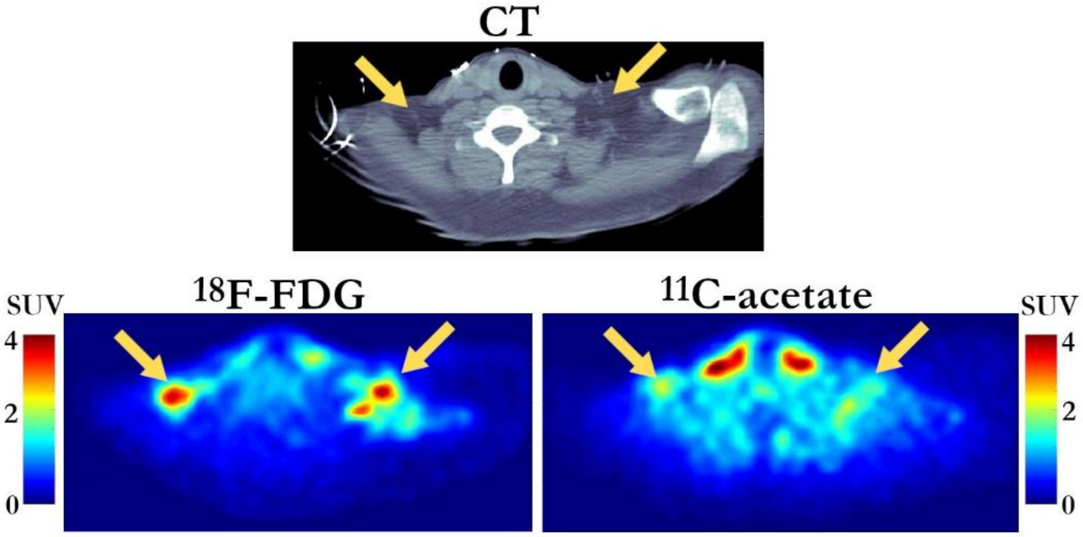


**Figure S4.** Typical positron emission tomography and computed tomography images of the shoulders and supraclavicular fat (arrows). $^{18}$F-FDG: fluorodeoxyglucose; CT: computed tomography; SUV: standardized uptake value. The $^{18}$F-FDG frame is the mean activity from t = 20-30 min (uptake phase). The $^{11}$C-acetate frame is the mean activity from t = 2-10 minutes (washout phase).

**Table S1. Baseline characteristics of study participants***

| *N* | 10 |
|---|---|
| **Age (years)** | 26.9 (22.9–31.0) |
| **Weight (kg)** | 83.4 (77.1–89.8) |
| **Height (m)** | 183.9 (180.9–186.9) |
| **BMI (kg/m$^2$)** | 24.7 (22.3–26.6) |
| **Fasting glucose (mM)** | 4.9 (4.6–5.2) |
| **Fasting insulin (pM)** | 58.7(44.4–72.9) |
| **Fasting FFA (mM)** | 0.32 (0.20–0.43) |
| **Fasting TG (mM)** | 0.90 (0.45–1.34) |
| **Fasting C-peptide (nM)** | 0.23 (0.16–0.30) |
| **Fasting leptin (ng/ml)** | 0.82 [0.34–1.88] |
| **Fasting glucagon (ng/l)** | 47.1 (27.2–67.0) |
| **Fasting GIP (pM)** | 8.8 (6.8–10.9) |
| **Fasting total GLP-1 (pM)** | 37.7 (29.9–45.4) |
| **HOMA-IR** | 1.83 (1.38–2.28) |

*Only participants who completed the study. Includes participant 13 who was excluded from analyses of carbohydrate-enriched diets (see Figure S1). Values are mean (95% confidence interval) for normally distributed data and median [interquartile range] for nonparametric data. BMI: body mass index; FFA: free fatty acids; GIP: glucose-dependent insulinotropic polypeptide; GLP-1: glucagon-like peptide-1; HOMA-IR: homeostatic model assessment for insulin resistance; TG: triglycerides.

**Table S2. MRI Parameters**

| | Shoulder anatomy | Liver anatomy | Shoulder/liver fat-water separation | Shoulder/liver fat-water separation |
|---|---|---|---|---|
| **Sequence type** | Turbo spin echo | Gradient echo | mDixon Quant | mDixon* |
| **Weighting** | T1 | T1 | T1 | T1 |
| **Echo train length** | 7 | 1 | 6 | 6 |
| **Echo time (ms)** | 16** | 1.4 | 0.97*** | 0.95*** |
| **Echo spacing (ms)** | 8 | | 0.7 | 1.4 |
| **Repetition time (ms)** | 542 | 3.04 | 5.67 | 9.04 |
| **Matrix size (voxel)** | 576 x 576 | 432 x 432 | 192 x 192 | 224 x 224 |
| **Field of view (mm)** | 400 x 400 | 400 x 400 | 400 x 400 | 400 x 400 |
| **Slice number/thickness (mm)** | 36/4 | 111/4 | 77/6 | 100/5 |
| **Flip angle (°)** | 90 | 10 | 3 | 5 |
| **Number of signal averages** | 4 | 1 | 1 | 1 |
| **Breath hold** | No | Yes | Yes | Yes |
| **View** | Axial | Axial | Axial | Axial |

Sequences were repeated for all MRI scans. * The commercial *mDixon Quant* sequence was not available for participant #1 and was replaced by an in-house sequence. The two Dixon sequences were performed for the remaining participants for comparison purposes. **Effective echo time. ***Time of first echo.

**Table S3. MRS Parameters**

| | BAT proton spectroscopy | Liver proton spectroscopy | scWAT proton spectroscopy |
|---|---|---|---|
| **Sequence type** | PRESS | STEAM | PRESS |
| **Echo train length** | 1 | 5 | 1 |
| **Echo time (ms)** | 30 | 10* | 30 |
| **Echo spacing (ms)** | | 5 | |
| **Number of signal averages** | 32 | 1 | 32 |
| **Repetition time (ms)** | 3000 | 3500 | 3000 |
| **Voxel size (mm)** | 8 × 8 × 8** | 25 × 25 × 25 | 10 × 10 × 10 |
| **Number of voxels** | 3 | 1 | 1 |
| **Spectral width (Hz)** | 2000 | 1250 | 2000 |

Sequences were repeated for all MRS scans. *Time of first echo. **Voxel size could be increased to 10 x 10 x 10 $mm^3$ when possible. BAT: brown adipose tissue; scWAT: subcutaneous white adipose tissue.
